# Enhanced Notch dependent gliogenesis and delayed physiological maturation underlie neurodevelopmental defects in Lowe syndrome

**DOI:** 10.1101/2024.11.25.625332

**Authors:** Yojet Sharma, Priyanka Bhatia, Gagana Rangappa, Sankhanil Saha, Padinjat Raghu

## Abstract

The activity of signaling pathways is required for coordinated cellular and physiological processes leading to normal development of brain structure and function. Mutations in *OCRL*, a phosphatidylinositol 4,5 bisphosphate [PIP_2_] 5-phosphatase leads to the neurodevelopmental disorder, Lowe Syndrome (LS). However, the mechanism by which mutations in *OCRL* leads to the brain phenotypes of LS is not understood. We find that on differentiation of LS patient derived iPSC, developing neural cultures show reduced excitability along with enhanced P levels of Glial Fibrillary Acidic Protein. Multiomic single-nucleus RNA and ATAC seq analysis of neural stem cells generated from LS patient iPSC revealed an enhanced number of cells with a gliogenic cell state. RNA seq analysis also revealed increased levels of *DLK1*, a non-canonical Notch ligand in LS patient NSC associated increased levels of cleaved Notch protein and elevation of its transcriptional target *HES5*, indicating upregulated Notch signaling. Treatment of iPSC derived brain organoids with an inhibitor of PIP5K, the lipid kinase that synthesizes PIP_2_, was able to restore neuronal excitability and rescue Notch signaling defects in LS patient derived organoid cultures. Overall, our results demonstrate a role for PIP_2_ dependent regulation of Notch signaling, cell fate specification and development of neuronal excitability regulated by OCRL activity.

## Introduction

The human brain is a complex organ whose development involves the regulated activity of many genes. In many cases, understanding how a gene controls brain development has come from studying brain disorders in human patients that harbor mutations in specific genes. These conditions, referred to as neurodevelopmental disorders (NDD) are a heterogeneous group of conditions in which individuals present with abnormalities of brain structure and/or function^1,2^. NDD represent the single largest class of monogenic inherited brain disease^3^ listed in Online Catalog of Human Genes and Genetic Disorders (OMIM^®^) database (https://omim.org/). Therefore, the study of cellular and developmental mechanisms underpinning NDD can lead to an understanding of normal brain development. Conversely, an understanding of underlying cellular and molecular mechanisms in NDD can help devise better strategies for the clinical management of patients.

The discovery of a diverse set of genes underlying NDD has led to a better understanding of cellular and development processes underpinning human brain development. Key examples include the identification of doublecortin (*DCX*) as the gene underpinning abnormal cerebral cortex development and morphology in patients with X-linked lissencephaly (XLS); analysis of XLS has underscored the importance of normal neuronal migration during cerebral cortex development^4,5^. Likewise, altered brain size and malformations in patients with mutations in the lipid phosphatase *PTEN*^6^ or the PI3K/AKT pathway^7,8^ [reviewed in^9^] has revealed the importance of lipid signaling in normal brain development. Mutations in *NOTCH2NL,* a human specific gene that regulates Notch signaling have underscored the importance of Notch regulated neurogenesis in the development of normal size and structure in the human neocortex^10^.

Lowe syndrome (LS) is a congenital X-linked recessive disorder in which three organs are affected, the eye, kidney and the brain^11^. Young boys with LS present with a cataract in early life, proximal renal tubular dysfunction and various degrees of neurodevelopmental deficit. The neurological features of LS include delayed cognitive milestones, ranging from mild intellectual disability to severe mental retardation, susceptibility to febrile seizures, hypotonia and in later years psychiatric symptoms. Imaging of brain structure in LS patients through Magnetic Resonance Imaging (MRI)^12,13^ reveals normal brain size with features of delayed myelination and focal, periventricular cystic lesions that develop with age; these findings are not specific to but consistent with gliotic changes. LS results from a mutation in the ***<u>O</u>***culo ***<u>C</u>***erebro ***<u>R</u>***enal syndrome of ***<u>L</u>***owe *OCRL* gene^14^. *OCRL* encodes for a protein that is part of a larger family of phosphoinositide 5 phosphatases^15^; the OCRL protein mainly shows phosphatase activity on phosphatidylinositol 4,5 bisphosphate (PIP_2_) generating phosphatidylinositol 4 phosphate (PI4P)^16^. OCRL localizes to a number of cellular organelles including the plasma membrane, Golgi, lysosomes and early endosomal compartments^17^ and multiple studies have implicated OCRL in the regulation of vesicular transport and the actin cytoskeleton [reviewed in^17,18^]. While many studies have attempted to address the function of *OCRL* in relation to the kidney phenotypes noted in human LS patients, much less is known about the cellular and developmental basis of the brain phenotypes in this disease. Some studies have sought to create models of LS with a view to understand brain phenotypes, including a zebrafish model^19–21^. It has been noted that a humanized mouse model for LS does not show phenotypes that recapitulate the cognitive impairment noted in LS patients although a hypotonia secondary to the renal manifestations has been reported^22^. This coupled with the inability to obtain biopsy samples from LS patient brains has severely impaired the understanding of the neurodevelopmental basis of LS syndrome.

In this study, we report the cellular, developmental and physiological phenotypes of 2D and 3D brain organoid cultures differentiated from LS patient derived induced pluripotent stem cell (iPSC) lines as well as an isogenic knockout of *OCRL* in wild type iPSC. We find that *OCRL* depleted, iPSC derived neurons (iNeurons) exhibit reduced neuronal excitability than controls. This phenotype is associated with an increase in the number of Glial Fibrillary Acidic Protein (GFAP) positive cells in LS patient cultures. Multiomic analysis of neural stem cells (NSC) derived from LS patients revealed an enhanced number of cells bearing molecular signatures of glioblasts and astrocytes associated with enhanced Notch signaling. These findings could be partially rescued by chemical inhibitors that reduce the synthesis of PIP_2_, the substrate of OCRL. Together, these findings suggest that during neural development, OCRL function is required to regulate PIP_2_ processes that control cell fate specification in the developing brain leading to an imbalance of neuron/astrocyte ratios, altered physiological development and brain function.

## Results

### *OCRL* is expressed during human brain development

To understand the role of *OCRL* in neurodevelopmental defects observed in Lowe Syndrome (LS) patients, we analyzed the gene expression pattern of *OCRL* in the developing human brain using a single-cell multiomics dataset ^23^ that included 38 biological samples divided into 27 brain regions from across five major developmental stages, ranging from the first trimester to adolescence. This dataset allowed us to map *OCRL* expression at the transcriptomic level across various cell types in the developing brain. A UMAP projection of this dataset reveals distinct clusters representing major cell types of the brain, including radial glia, intermediate progenitor cells (IPC), neurons, astrocytes, oligodendrocytes and microglia (Fig 1A). In addition, *OCRL* mRNA was expressed at varying levels depending on the developmental stage and cell type. The dot plot shows that OCRL expression was detected throughout development, with the highest expression detected during the 246-498 days post conception (Fig 1B). Further, analysis of cell-type specific enrichment of *OCRL*, revealed that in this dataset, this gene was predominantly expressed in GABAergic and glutamatergic neurons compared to other cell-types (Fig 1C). These observations suggest that *OCRL* plays a role during *in utero* brain development.

**Figure 1:**
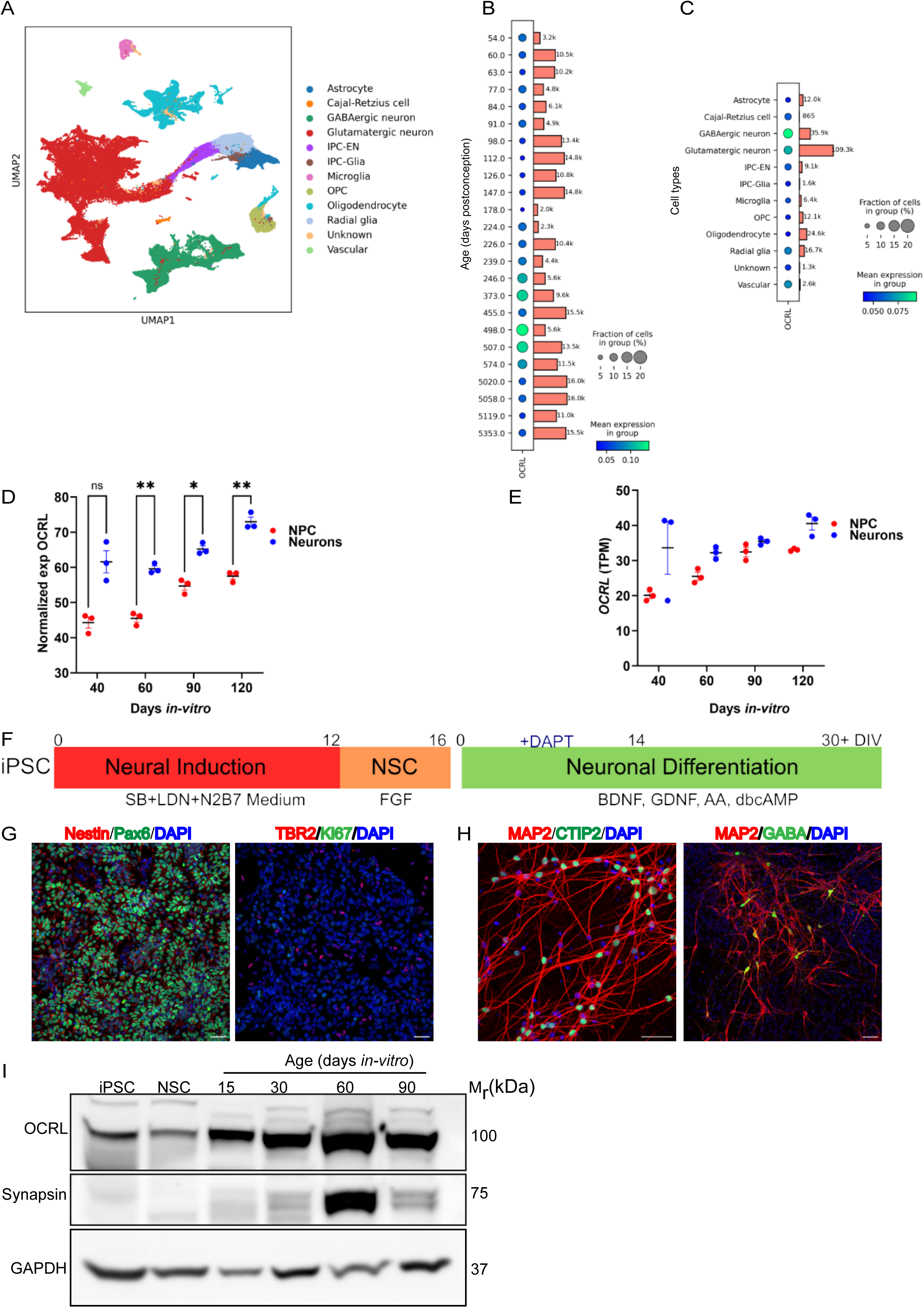
OCRL expression during human brain development. **(A)** UMAP plot derived from a single nuclei multiome dataset of Wang et.al ^23^, illustrating the various cell types in the developing human brain. These include Astrocytes, Cajal-Retzius cells, GABAergic neurons, Intermediate Progenitor Cells (IPC) for Excitatory Neurons (IPC-ENs), IPCs for Glia (IPC-Glia), and Radial Glia cell clusters. **(B)** Dot plot showing the expression of *OCRL* at distinct time points during human brain development. Y-axis is age in days post-conception. Size of the dots represent the proportion of cells expressing *OCRL*. The colour gradient represents the mean expression in each group of cells. Bars represent the number of cells for a given time point in this dataset. **(C)** Dot plot displaying *OCRL* expression across distinct cell types present in this dataset. **(D)** Mass spectrometry-based proteome analysis of SOX2::EGFP+ neural stemstem cells (NSC, red) and hSYN1::dTomato+ neurons (blue), showing the expression of OCRL at different stages of development in iPSC-derived human brain organoids. Statistical significance was determined by an unpaired t-test with Welch’s correction. **(E)** Bulk RNA sequencing of SOX2::EGFP+ neural stem cells (NSC, red) and hSYN1::dTomato+ neurons (blue).Y-axis shows transcript levels in TPM (specific transcript counts per million total counts. X-axis show age of culture as days *in vitro* revealing the temporal expression patterns of OCRL during different stages of development in iPSC-derived human brain organoids. **(F)** Schematic representation of the Dual-SMAD inhibition protocol used for generating dorsal forebrain neural stem cells (NSC). iPSC-induced pluripotent stem cells, Neural Induction (conversion of iPSC into NSC; Neural differentiation (conversion of NSC into neurons); N2-alternative supplement for culturing NSC without serum, B27 (supplement of antioxidant enzymes, proteins, vitamins, and fatty acids), BDNF-brain derived neurotrophic factor; GDNF-Glia derived neurotrophic factor, DAPT-N-[N-(3, 5-difluorophenacetyl)-l-alanyl]-s-phenylglycinet-butyl ester, dbcAMP – dibutryl cyclic adenosine monophosphate, AA-Ascorbic Acid. **(G)** Immunofluorescence images of WT1-derived NSC, showing expression of Nestin (red) and Pax6/Foxg1 (green); Ki67 (green) and TBR2 (red) expression is also shown. **(H)** NSC terminally differentiated into mature neurons were identified by MAP2 (red) and CTIP2 (green, a layer 6 neuron marker). The cultures were further stained to identify GABAergic interneurons (GABA, green) and MAP2. Nuclei were counterstained with DAPI (blue). Scale bar = 50 µm. **(I)** Western blot analysis of OCRL expression in the WT1 cell line, showing OCRL protein levels in iPSCs, NSCs and during 15, 30, 60, and 90 days *in vitro (*DIV) neuronal cultures. Levels of synapsin 1, a mature neuronal marker are shown. GAPDH was used as a loading control. Samples from iPSC, NSC and neuronal cultures of different ages differentiated from NSC are shown.

We also examined expression of the OCRL protein in iPSC-derived human brain organoids at different stages of *in vitro* development using a publicly available proteomics dataset ^24^. This revealed that at all stages of *in vitro* differentiation, hSyn+ mature neurons expressed higher levels of OCRL compared to Sox2+ NSC (Fig 1D). Likewise in a transcriptomic data set, *OCRL* transcript was seen in both neurons and NSC (Fig 1E). Importantly, OCRL expression increased progressively over time, indicating a potential role in neuronal maturation and further supporting the age-specific enrichment observed in foetal brain samples (cf. Fig 1B).

To validate these findings, we utilized a previously established wild-type control iPSC line (WT1)^25^ to generate NSC using the dual-SMAD inhibition protocol (Fig.1F, G).Upon terminal differentiation into cortical neurons, cells expressed mature neuronal markers such as MAP2 (pan-neuronal), CTIP2 (deep-layer), and GABA (interneurons) (Fig 1H). Western blot analysis of OCRL expression at various time points (iPSC, NSC, 15, 30, 60, 90DIV) confirmed a gradual increase in OCRL protein levels as differentiation progressed, with higher levels in mature neurons compared to NSC or early-stage neurons (Fig 1I). Overall our findings support an age dependent increase in OCRL protein expression during neuronal differentiation.

### Loss of *OCRL* leads to reduced neuronal activity during development

We differentiated iPSC derived from three individual patients (LSP) with LS from a single family^26^ ^27^. These include two monozygotic twins (LSP2 and LSP3) and their maternal cousin (LSP4). These iPSCs were differentiated into NSC which expressed the characteristic protein markers Nestin, Pax6 and FoxG1 (Supp. Fig 1A). Further terminal differentiation into cortical neurons was confirmed by the expression of mature neuronal markers MAP2 and CTIP2 (Fig 2A).

**Figure 2:**
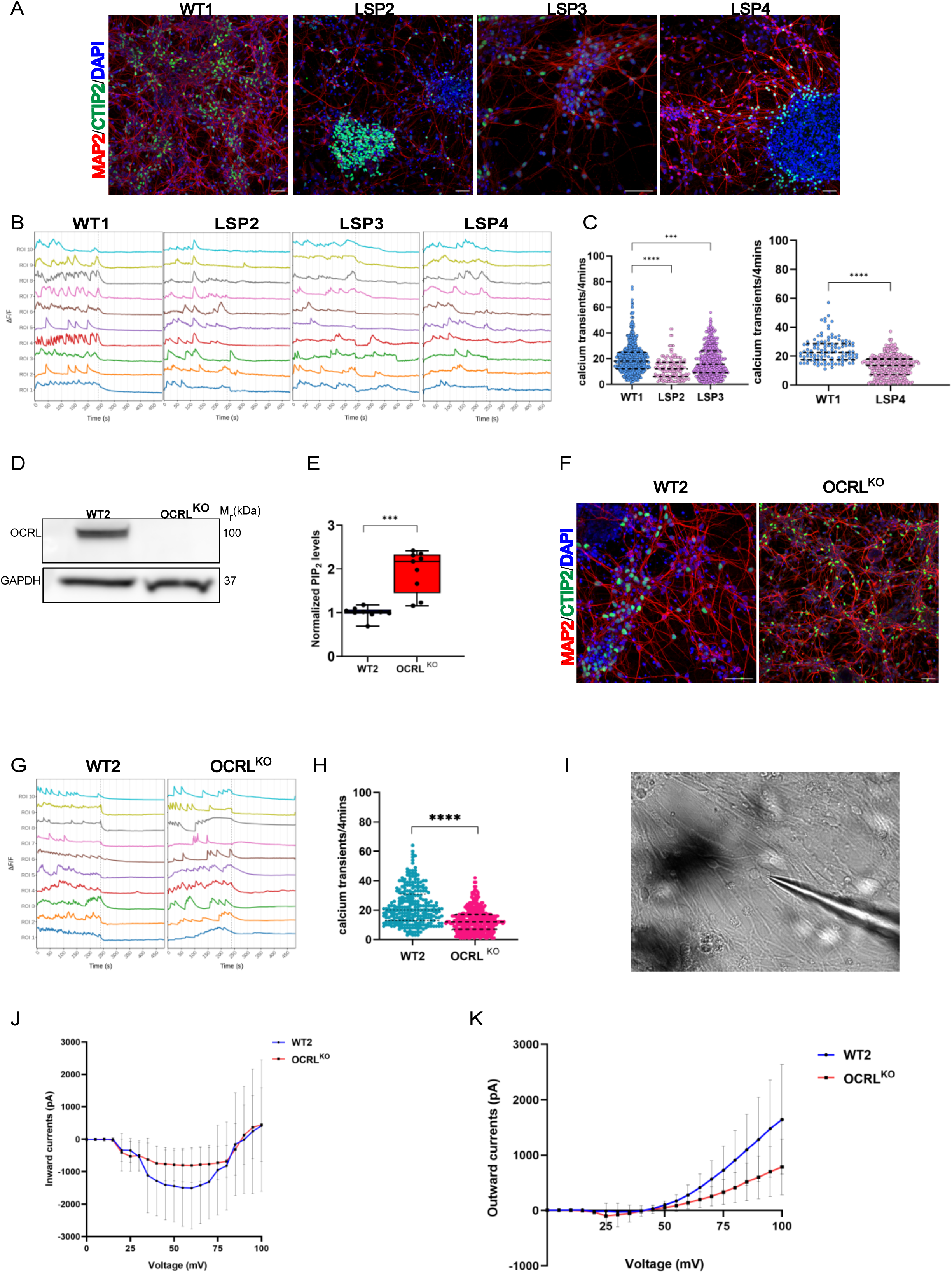
Altered physiological activity in OCRL depleted developing neurons. **(A)** Immunofluorescence images of 30-day *in-vitro* (DIV) neurons from WT1, LSP2, LSP3, LSP4, stained for mature forebrain cortical neuron markers MAP2 (red) and CTIP2 (green). Nuclei are stained with DAPI (blue) **(B)** Representative calcium ([Ca²⁺]i) transient traces from 30 DIV neurons of WT1, LSP2, LSP3, and LSP4, normalized to the first frame. Each trace (of a different colour, represents [Ca²⁺]i transients from the soma of a single neuron (ROI). Y-axis represents change in represents [Ca²⁺]i shown as ΔF/F. A dotted line marks the addition of TTX (1 µM), which abolishes calcium transients **(C)** Scatter plots showing the frequency of calcium transients per 4 minutes for WT1, LSP2, LSP3 and LSP4. Unpaired t-tests with Welch’s correction were used to analyze data from 150-600 neurons across multiple differentiations. The WT1 and LSP4 data include 115-270 neurons. **(D)** Western blot showing OCRL protein expression in WT2 and OCRL^KO^ neurons, with GAPDH as the loading control. **(E)** Liquid chromatography-mass spectrometry (LC-MS) analysis measuring total PIP₂ levels in 30 DIV WT2 and OCRL^KO^ neurons. **(F)** Immunofluorescence images of 30 DIV neurons from WT2 and OCRL^KO^ stained for mature forebrain cortical neuron markers MAP2 (red) and CTIP2 (green). Nuclei are stained with DAPI (blue) **(G)** Representative calcium transient traces for 30 DIV WT2 and OCRL^KO^ neurons. **(H)** Frequency of [Ca²⁺]_i_ transients in 150-275 neurons from WT2 and OCRL^KO^. **(I)** Showing a representative image of patched 60 DIV neuron. Whole-cell patch clamp recording of WT2 and OCRL^KO^ neurons in voltage-clamp mode exhibits **(J)** inward currents and **(K)** outward currents respectively. Y-axis -current in pA; X-axis voltage in mV. Error bars represent Mean+/- SD for currents recorded form multiple cells at each voltage.

To measure neuronal excitability, we performed intracellular calcium ([Ca^2+^]_i_) imaging on both WT1 and LSP neurons (LSP2, LSP3, LSP4). Representative [Ca^2+^]_i_ traces from each genotype are shown in Fig 2B. Quantification revealed a significant reduction in the frequency of [Ca^2+^]_i_ transients in all three LSP neurons compared to WT1 controls (Fig 2C). To test the dependence of this phenotype noted in LSP derived neurons on OCRL function, we generated a knockout iPSC line (OCRL^KO^) using CRISPR-Cas9 genome engineering; a stop codon was introduced in exon 8 thus recapitulating the *OCRL* allele seen in the LSP used in this study (Suppl Fig1B-E). As an isogenic control (WT2), we used the parent line (NCRM5) that was used to generate OCRL^KO^. The WT2 and OCRL^KO^ iPSC were positive for pluripotency markers SSEA4, SOX2, TRA-160 and OCT4 (Suppl Fig1D) and OCRL^KO^ iPSC also had normal karyotype (Suppl Fig1E). We generated NSC from WT2 and OCRL^KO^ (Suppl Fig 1F) and differentiated these into neurons (Fig 2F). Western blot analysis confirmed the absence of OCRL protein in OCRL^KO^ neurons compared to WT2 controls (Fig 2D). To test the biochemical consequence of OCRL depletion in OCRL^KO^, we measured the levels of PIP_2_ from 30 DIV neurons using liquid chromatography, mass spectrometry. As expected, PIP_2_ levels were significantly elevated in OCRL^KO^ neurons relative to WT2 controls (Fig 2E), recapitulating the findings previously reported by us for LSP lines^27^. [Ca^2+^]_i_ imaging on OCRL^KO^ neurons (Fig 2G) revealed a reduction in the frequency of [Ca^2+^]_i_ transients in OCRL^KO^ compared to WT2 (Fig 2G, H) recapitulating our findings in LSP derived neurons (Fig 2B,C).

To corroborate our finding of reduced neuronal excitability in OCRL depleted neurons, we performed whole-cell patch-clamp recordings on 60DIV neurons. Resting membrane potential (Suppl Fig 2A) and capacitance measurements (Suppl Fig 2B) were no different between WT2 and OCRL^KO^. However inward currents were reduced in OCRL^KO^ neurons compared to WT2 (Fig 2J) and outward currents also showed a similar trend of being reduced in OCRL^KO^ (Fig 2K). Similar patterns were observed in recordings from LSP2 and LSP3 patient-derived neurons compared to WT1 at 60 DIV, particularly for inward currents. However, we noted that fewer patient-derived neurons were available for recording at this time point, limiting our sample size (Suppl Fig 2C, D).

### *OCRL* depletion alters cell type composition in iPSC derived neural cultures

During this study we stained iPSC derived cultures from control and OCRL depleted lines for markers of neurons and astrocytes. Cultures stained for the mature astrocytic marker GFAP, showed an increase in staining in each of the three LS patient lines compared to WT1 at 30 DIV (Fig 3A). This increase in GFAP staining was confirmed by Western blot analysis, which showed elevated GFAP protein expression in LSP neural cultures relative to WT1 (Fig 3C), with quantification revealing an increased fold change in each of the LSP lines compared to WT1 (Fig 3D). By contrast staining with S100β (immature marker for astrocytes) revealed fewer numbers of S100β cells in LS patient lines compared to WT1 (Fig 3A). In the same cultures, there was no change in the staining pattern of the neuronal markers MAP2 and CTIP2 (Fig 3B). To test if depletion of OCRL was sufficient to produce this phenotype, we compared cultures differentiated from WT2 with OCRL^KO^ at 30DIV. Immunofluorescence images from 30DIV OCRL^KO^ neural cultures also displayed an increased number of GFAP+ cells compared to their isogenic WT2 controls (Fig 3E) and western blot analysis confirmed significantly higher GFAP expression observed in OCRL^KO^ neural cultures compared to WT2 (Fig 3F, G).

**Figure 3:**
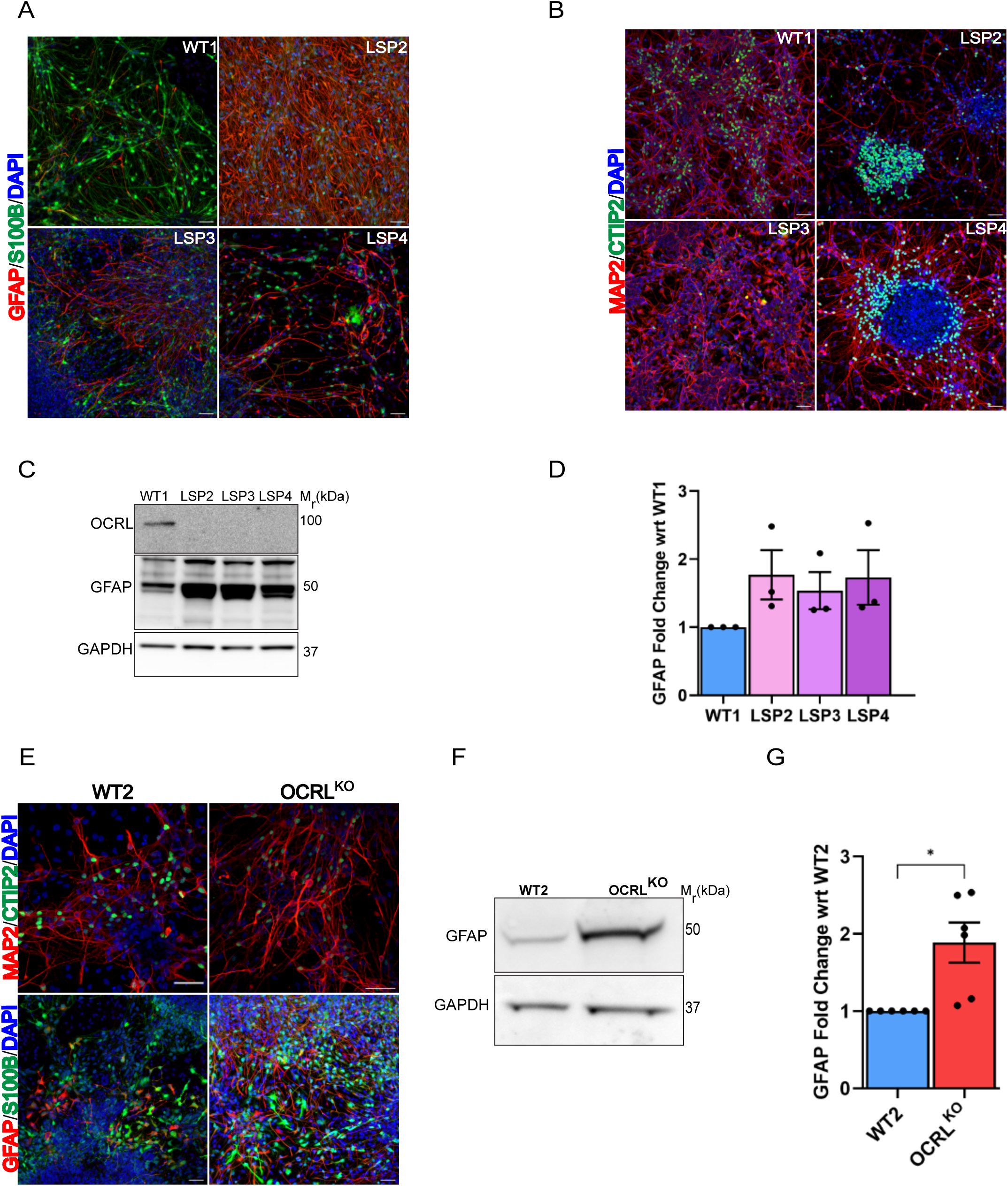
Altered cellular composition on OCRL depletion. 30 DIV neural cultures from WT1, LSP2, LSP3, LSP4 stained with **(A)** mature astrocytic marker GFAP (red) and immature astrocytic marker (green) S100β respectively, **(B)** MAP2 (red) and CTIP2 (green). Scale-bar=50μm. **(C)** Western blot with protein lysates from 30DIV iPSC derived cultures of WT1, LSP2, LSP3, LSP4 cell lysates; immunoblotting for OCRL and GFAP protein are shown; GAPDH 37kDa was used as a loading control. **(D)** GFAP fold-change in LSP derived 30 DIV neural cultures analysed and plotted w.r.t the control WT1 across three independent neural inductions. Y-axis shows the fold change in GFAP levels in LSP relative to WT1 **(E)** 30 DIV WT2 and OCRL^KO^ neurons were also stained with GFAP (red) and S100β (green), MAP2 and CTIP2. Scale bar=50μm. **(F)** Western blot of GFAP for WT2 and OCRL^KO^ showing levels of GFAP protein in lysates of 30 DIV neurons**. (G)** GFAP fold-change in GFAP levels analysed and plotted w.r.t WT2 control across five independent neural inductions. Unpaired t-test with Welch’s correction was used to calculate statistical significance.

### Altered cellular neurodevelopmental state in Lowe syndrome patient NSC

Since we noted an increased expression of the glial marker GFAP in LSP and OCRL^KO^ cultures on terminal differentiation of NSC into neural cultures, we wondered if OCRL depleted NSC had altered cell states compared to controls. To test this, we carried out single nuclei analysis of the NSC population derived from iPSC of WT1 and the LSP lines LSP2 and LSP3. For our experiments, we employed single-nuclei multiomics (scMultiome) analysis, which integrates both transcriptomic (RNA-seq) and chromatin accessibility (ATAC-seq) and allows the simultaneous assessment of gene expression and chromatin accessibility at single-cell resolution^28,29^.

The workflow for our scMultiome analysis is outlined in Suppl Fig 3A. Nuclei were isolated from LS patient-derived NSCs (LSP2, LSP3) and wild-type controls (WT1), followed by transcriptome (RNA-seq) and chromatin accessibility (ATAC-seq) data generation using the 10X Chromium Single Nuclei Multiome ATAC + Gene Expression kit. After pre-processing and quality control, we retained 19780 nuclei for WT1, 19854 nuclei for LSP2 and 19953 nuclei for LSP3 for analysis. The UMAP generated from the single cell RNA-seq data showed three distinct clusters corresponding to WT1, LSP2 and LSP3 (Fig 4A). Using the Leiden graph-based clustering method, we obtained 14 clusters (Fig. S3B). Each cluster was defined by a set of highly expressed transcripts (Suppl. Fig 3F). To ensure robust cluster annotation, we employed two levels of annotation: a first level was performed using snapseed^30^ and a second level of cluster annotation was carried out using scVI-scanVI framework^31^ to refine cluster identities into more specific cell-states. To this end, we leveraged already available snRNA/snMultiome reference of developing human brain datasets ^32^ ^23^ for accurate mapping of cell states using the scVI-scanVI framework (Suppl Fig.3C) First, using snapseed, we obtained preliminary marker-based hierarchical cell type annotation. Second, we projected our NSC dataset to the latent space of the developing human reference datasets using scVI and scanVI, and generated confusion matrices (Suppl. Fig3 D, E). Cell type labels were assigned based on the highest confidence scores, represented by the yellow and green bands in the confusion matrices (Suppl. Fig3 D, E). These high confidence matches in the transfer learning approach were used to establish the final cell type annotations, resulting in the labeled UMAP visualization (Fig 4B). Notably, LSP2 NSCs showed 0.5% (112 nuclei) corresponding to astrocytes, 86% (17096 nuclei) to glioblast. On the other hand, LSP3 NSCs had 28% nuclei (5730) for astrocytes and 57.53% towards glioblast cells. WT1 NSCs had 54.91% (10862) radial-glia cells, but only 0.16% (33) for glioblast and none for astrocytes (Fig. 4C). This marked increase in glioblast and astrocyte cell states compared to wild-type controls is consistent with the enhanced number of GFAP positive cells previously observed in LS patient-derived neural cultures (Fig 3A).

**Figure 4:**
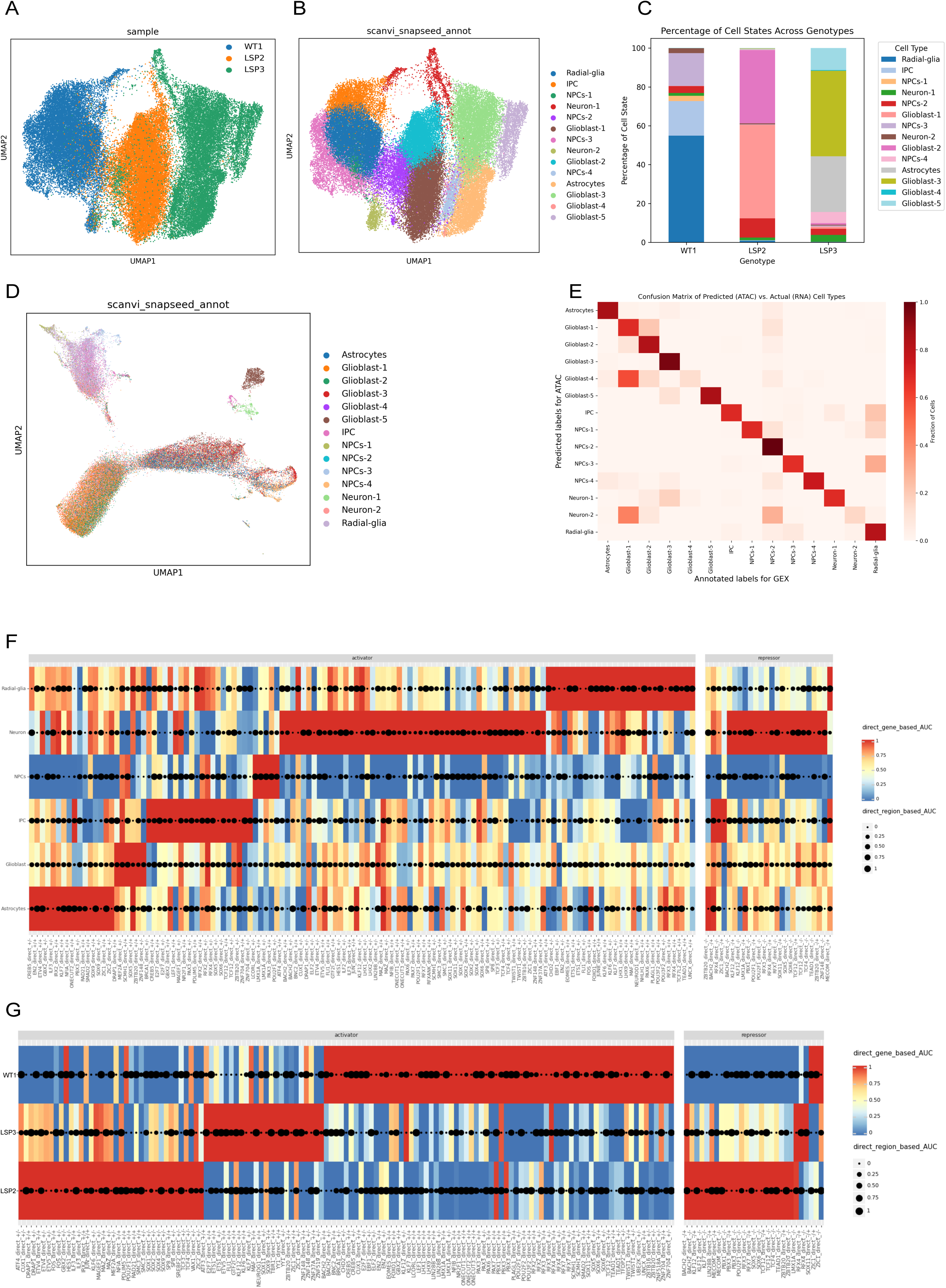
Single cell analysis of OCRL depleted NSC. **(A)** UMAP projection of single-nuclei RNA-seq (snRNA-seq) data from WT1, LSP2, and LSP3 samples, showing the distribution of cells across the three genotypes. **(B)** UMAP projection of snRNA-seq data with cell types annotated using scVI-scANVI mapping. Major cell types include astrocytes, glioblasts, intermediate progenitor cells (IPCs), neuronal stem cells (NSC 1-4), neurons (1-2), and radial glia are shown. **(C)** Stacked bar plot showing the percentage of each cell type in WT1, LSP2, and LSP3 samples. X-Axis shows the samples identity; Y-axis shows the % of cells of each cluster per sample**. (D)** UMAP projection of ATAC-seq data from the cells used in the multiome experiment, coloured by cell types defined in panel B analysed using snapATAC2. **(E)** Confusion matrix comparing cell-type annotations between ATAC-seq and gene expression (GEX) data. X-axis shows annotated labels for GEX and Y-axis shows predicted labels for ATAC-seq. **(F)** Heatmap and dot plot representing transcriptional regulatory relationships across different cell clusters. The y-axis shows various cell clusters observed: radial-glia, neuron, NSCs, IPC, glioblast and astrocytes. The x-axis shows transcription factors (lower axis) and divided into two categories: activators and repressors (upper axis). The symbol between brackets indicates whether the TF activates (+) or represses (-) its target genes. The colour intensity represents direct gene-based Area Under the Curve (AUC) values (ranging from 0 (blue) to 1 (red), indicating the degree of enrichment or specificity of transcription factors in each cell type. The size of black dots represents direct region-based AUC values (ranging from 0 to 1), indicating how well a transcription factor’s binding sites are enriched in regulatory regions specific in each cell type. Larger dots indicate stronger enrichment of the transcription factor at regulatory regions characteristic of a cell type. **(G)** Heatmap and dot plot representing transcriptional regulatory relationships across different samples. The y-axis shows sample clusters: WT1, LSP2, LSP3. The x-axis shows transcription factors (below heat map) divided into two categories: activators and repressors (above heat map). The symbol between brackets indicates whether the TF activates (+) or represses (-) its target genes. The colour intensity represents direct gene-based Area Under the Curve (AUC) values (ranging from 0 to 1, blue to red), indicating the degree of enrichment or specificity of transcription factors in each cell type. The size of black dots represents direct region-based AUC values (ranging from 0 to 1), indicating how well a transcription factor’s binding sites are enriched in regulatory regions specific to each sample type. Larger dots indicate stronger enrichment of the transcription factor at regulatory regions characteristic of that sample type.

To independently study the cell composition of LSP NSC, we used SnapATAC2 for clustering cells based on chromatin accessibility profiles. UMAP plots of the ATAC-seq data showed cells from WT1, LSP2 and LSP3 clustering separately (Suppl Fig 3G) with LSP2 and LSP3 clustering closer to each other and away from WT1. Further clustering led to the identification of clusters of cells corresponding to the 14 types (Fig 4D) (Suppl Fig 3H) also identified in the single nuclei RNA-seq data (Fig 4B). We found that an integration of transcriptome (RNA-seq) and chromatin accessibility (ATAC-seq) data showed strong concordance with each other, as evidenced by the diagonal pattern of high correlation (dark red) in the confusion matrix (Fig 4E).

To identify enhancer driven gene regulatory interactions, we applied SCENIC+^33^, a tool that identifies putative enhancer regions in the genome and their associated transcription factors, while simultaneously mapping these regulatory elements to their potential downstream target Motif analysis revealed enrichment of transcription factors (Suppl Table 1) associated with astrocyte lineage commitment, including nuclear factor one (NFI) family of transcription factors NFIA, NFIB, NFIC and NFIX, and SOX2, POU2F1 (OCT1) in glioblasts^34–36^. Additionally, radial glia-specific motifs such as LHX9 and EMX1 were identified in radial glia populations. Using, SCENIC+ we analyzed the multiome data in relation to cell type clusters asking which transcription factors, both activators and repressors were active in each cell type. We analyzed both the degree of enrichment and specificity of transcription factors as well as how well a transcription factor’s binding sites are enriched in regulatory regions specific to each cell type. This analysis revealed a signature of transcription factor activity for each cell type cluster (Fig 4F). For example, transcription factors and sites of active chromatin in neurons showed presence of neuron-specific transcription factors such as ONECUT1/3 that were not as enriched in glioblasts and astrocytes (Fig 4F)^37,38^. Simultaneously, we also generated the heatmap-dot plot of regulons sample-wise to understand the state of transcription factors of cells in each sample WT1, LSP2 and LSP3 (Fig 4G). This analysis revealed that WT1 was enriched for binding sites for transcription factors overlapping with those enriched in radial glial cells, intermediate progenitors and neurons (Fig 4G). By contrast, transcription factors enriched in LSP2 and LSP3 overlapped considerably with those identified for glioblasts and astrocytes such as ETV4^39^, SOX9 and NFIA (Fig 4H). Overall, our findings demonstrate that OCRL deficiency alters the transcriptional landscape of NSC, priming the LSP NSC towards accelerated gliogenesis.

### Depletion of *OCRL* leads to altered Notch signaling

During pseudobulk analysis of our single nuclear RNAseq dataset we noted that levels of the atypical Notch activator, DLK1^40,41^, were elevated in a large fraction of LSP2 and LSP3 NSC along with the long non-coding mRNA *MEG3* and *MEG8* that are found in the *DLK1-DiO3* locus (Fig 5A). We tested the status of Notch signaling in OCRL depleted cultures. Given the paracrine nature of Notch signaling that depends on cell-cell interactions we performed these assays with iPSC differentiation into 3D organoid cultures. WT1 and LSP iPSC were differentiated into organoids (Fig 5B, Suppl Fig 4A). These 3D organoid cultures recapitulated the phenotypes previously described using the 2D culture system (cf. Fig 3). RT-PCR analysis of 90 DIV organoids showed no change in the levels of the neuronal transcripts *MAP2* (Suppl Fig 4C) while the levels of astrocytic transcripts *NF1A* and *GFAP* was elevated (Suppl Fig 4D, E). Immunohistochemical analysis of 90 DIV organoid sections showed increased GFAP staining in LSP2, LSP3 and LSP4 compared to WT1 (Suppl Fig 4F). Western blot analysis from 25 DIV cultures revealed elevated levels of DLK1 protein in lysates from LSP2, LSP3 and LSP4 organoids (Fig 5C, D), and a similar elevation of secreted DLK1 was seen in supernatants from these cultures (Fig 5F). To test for Notch receptor activation, we measured the levels of the cleaved Notch receptor (cNotch) by Western blots; this revelated elevated levels of cNotch in LSP organoids compared to WT1 (Fig 5C, E) organoids. Lastly, we measured the levels of the Notch signaling dependent transcript *HES5* ^42^ and found it to be elevated in LSP2 and LSP3 relative to WT1 (Fig 5G). These observations of elevated DLKI protein levels (Fig H, I), cNotch (Fig 5H, J) and *HES5* transcript (Fig 5K) were also seen in OCRL^KO^ compared to WT2.

**Figure 5:**
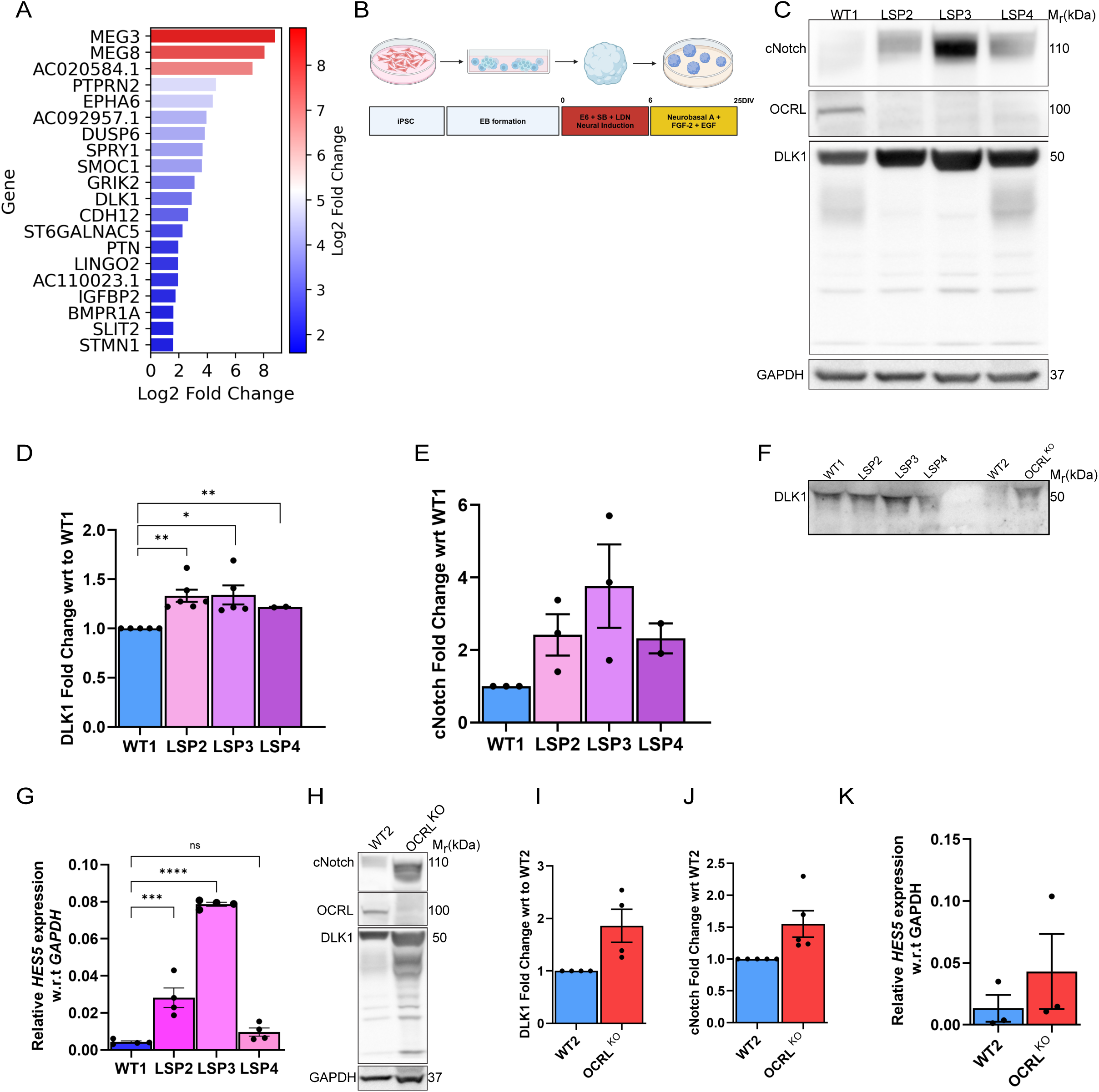
Altered Notch signaling in OCRL depleted organoid cultures. **(A)** Bar plot showing the top differentially expressed genes (DEGs) identified through pseudobulk analysis of single nuclei RNA-seq data between aggregated LSP (LSP2 and LSP3) and WT1 samples. The x-axis displays the Log2 Fold Change values, while the y-axis lists the top 20 enriched genes in LSP samples that satisfied a stringent significance criterion of log2 fold change > 1.5 and adjusted p-value < 0.01. A color gradient from blue (lower fold changes) to red (higher fold changes) indicates the magnitude of differential gene expression. **(B)** Schematic of the protocol for generation of 3D spheroids from human iPSC of controls and Lowe syndrome patients. SMAD inhibition [treatment with SB-31240 (10 μM) and LDN (100 nM)]. FGF-2 Fibroblast Growth Factor 2; EGF-Epidermal growth factor **(C)** Western blot with WT1, LSP2, LSP3 and LSP4 lysates obtained from 25 DIV brain organoids; detection of OCRL (100kDa), cleaved notch (cNotch, 110kDa), DLK1 (50kDa) is shown; GAPDH (37kDa) was used as a loading control. Quantification of **(D)** DLK1 levels and **(E)** cNotch levels in LSP lysates relative to WT1; Each point represents 10-15 brain organoid from independent batches of differentiation. Statistics: Unpaired student t-test using Welch correction. Error bars represent Mean +/- SEM **(F)** Western blot for secreted DLK1 (sDLK1) obtained from medium of WT1, LSP2, LSP3, LSP4, WT2, and OCRL^KO^ brain organoid cultures maintained till 25 DIV. Equal amount (40μg) of protein has been loaded per sample. **(G)** RT-PCR analysis for 25 DIV brain organoids from WT1, LSP2, LSP3, LSP4 for the cNotch downstream effector *HES5* normalized to the housekeeping gene *GAPDH*. Each dot represents 15-20 brain organoids generated from independent batches of differentiation. Mean +/- SEM shown. Statistical test used: Ordinary one-way ANOVA **(H)** Western blot from WT2 and OCRL^KO^ 25 DIV brain organoids for DLK1 and cNotch. GAPDH was used as a loading control. Quantification of DLK1 **(I)** and cNotch **(J)** bands for WT2 and OCRL^KO^ brain organoids. Y-axis represents protein levels in OCRL^KO^ relative to WT2 **(K)** *HES5* transcripts relative to the housekeeping gene GAPDH quantified and plotted for WT2 and OCRL^KO^ brain organoids. Error bars represent Mean +/- SEM.

### Elevated PIP_2_ levels underpin cell fate composition and neuronal excitability in OCRL depleted cultures

As an enzyme, OCRL dephosphorylates PIP_2_ to generate PI4P and OCRL depleted cells show elevation of PIP_2_ levels (Fig 2E). To test if this elevation of PIP_2_ levels is important for the phenotypes we report here, we pharmacologically inhibited the enzyme, phosphatidylinositol 4 phosphate 5-Kinase (PIP5K) that is responsible for synthesizing the majority of PIP_2_; the human genome PIP5K is encoded by three genes PIP5KA, PIP5KB and PIP5KC^43^. We differentiated WT2 and OCRL^KO^ NSC into neurons and treated the cultures for 7 days with an inhibitor for PIP5KC (UNC3230)^44^ (hereafter called PIP5K*1C_i_*) prior to analysis (Fig 6A). Following treatment with PIP5K1C_i_, the frequency of [Ca^2+^]_i_ transients in OCRL^KO^ neurons was elevated up to that seen in WT2 derived cultures (Fig 6B). Likewise, using the 3D organoid culture system, we tested the effect of treatment with PIP5K1C_i_ (Fig 6C) on the levels of DLK1 protein; these were reduced in OCRL^KO^ treated with PIP5K*1C_i_* compared to untreated OCRL^KO^ cultures (Fig 6 D, E). Likewise, the elevated levels of cNotch in OCRL^KO^ organoids were reduced on treatment with PIP5K1C_i_ (Fig 6 D, F). Lastly the elevated levels of *HES5* transcript in OCRL^KO^ was reverted to WT2 levels on treatment with PIP5K inhibitor (Fig 6G).

**Figure 6:**
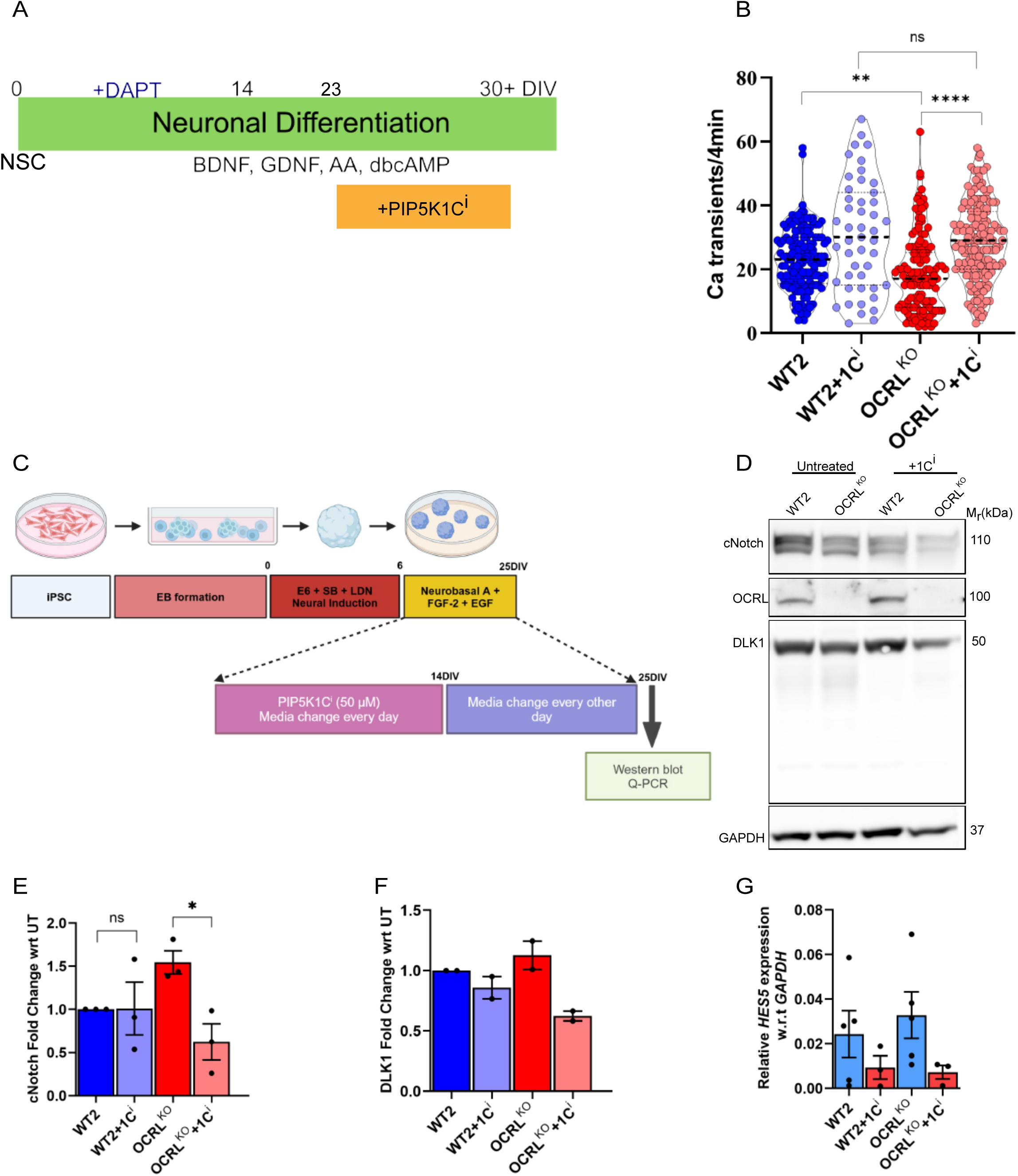
Rescue of phenotypes in OCRL depleted cultures by inhibiting PIP_2_ synthesis. **(A)** Schematic of experimental design for testing the effect of PIP5KC inhibitor on [Ca^2+^]_i_ transient frequency in OCRL depleted neurons. BDNF-brain derived neurotrophic factor; GDNF-Glia derived neurotrophic factor, DAPT-N-[N-(3, 5-difluorophenacetyl)-l-alanyl]-s-phenylglycinet-butyl ester, dbcAMP – dibutryl cyclic adenosine monophosphate, AA-Ascorbic Acid; PIP5KC_i_ – inhibitor of PIP5K (UNC-3230). **(B)** Quantification of [Ca^2+^]_i_ transients in 30 DIV WT2, OCRL^KO^ neurons and WT2 and OCRL^KO^ treated with PIP5K1C_i_ (10uM) for a duration of 7 days *in-vitro*. Y-axis represents the frequency of [Ca^2+^]_i_ transients **(C)** Schematic of experimental design for testing the effect of PIP5KC_i_ on Notch signaling **(D)** Western blot showing levels of cNotch and DLK1 in 25DIV organoids from WT2, OCRL^KO^ (untreated), WT2, OCRL^KO^ treated with PIP5K1C_i_. Quantification of cNotch **(E)** and DLK1 **(F)** protein levels presented as fold-change w.r.t to untreated condition respectively. **(G)** Transcript levels of *HES5* measured using Q-RTPCR for WT2, OCRL^KO^ 25DIV organoids treated with PIP5K1C_i_ presented relative to *GAPDH* transcripts. Each point shown is a single RNA sample from 15-20 organoids: 5 replicates. Error bars: Mean +/- SEM.

## Discussion

During brain development, differentiating neurons progressively develop activity that can be monitored in experimental system as [Ca^2+^]_i_ transients; these transients have been shown to be critical for a number of neurodevelopmental events such neurite outgrowth and neurotransmitter specification [reviewed in^45^]. In the case of LS, the cellular and physiological changes in the developing brain that manifest with clinical features of altered higher mental function are not known. In this study, using several independent LS iPSC derived disease-in-a-dish model, we found that neuronal cultures derived from LS patients showed reduced numbers of [Ca^2+^]_i_ transients and inward currents underscoring the reduced neuronal activity. Thus, it is likely that as the brain of LS patients develops *in utero,* neuronal activity is also reduced.

We found that the reduced neuronal activity in LS patient derived cultures could be recapitulated when wild type iPSC was edited to introduce a specific mutation in exon 8 of *OCRL* that truncates the protein prior to the phosphatase domain, equivalent to the mutant allele in the LS patient lines we have studied. These findings underscore the critical role of OCRL in the development of neuronal activity in the brain. OCRL is a PI(4,5)P_2_ phosphatase and levels of this lipid are elevated when the enzyme activity is lost. We found that (i) the levels of PIP_2_ are elevated in LS patient cells^27^ ^46^ and in OCRL^KO^ (Fig 2-this study) (ii) The reduced neuronal activity could be rescued by treating OCRL^KO^ cells with an inhibitor of PIP5KC enzyme that synthesizes PIP_2_. Taken together, these findings strongly suggest that the elevated PIP_2_ levels lead to the reduced neuronal activity during brain development. PIP_2_ regulates several key molecules involved in regulating neuronal excitability including ion channels and transporters^47,48^ and also synaptic vesicle cycle proteins^49^. Altered function of these molecules could underlie the reduced excitability observed in developing OCRL depleted neurons.

Brain development can be conceptualized as a series of cell specification events followed by differentiation into neural cell types that wire together forming circuits and develop activity and function. At which stage of this process is OCRL required for normal activity development? During our experiments, we noted that neural cultures derived from OCRL depleted iPSC expressed higher levels of GFAP that marks mature astrocytes. Increased numbers of astrocytes have previously been noted in other neurodevelopmental disorders including autism^50^ and Down’s syndrome^51^. While there are no studies on the histology of the developing brain from LS patients to directly corroborate our finding in human patients, a previous analysis of LS using the zebrafish model have revealed features suggestive of increased focal gliosis when OCRL is depleted^19^. This finding of increased astroglia in LS patient derived neural cultures raises the possibility these astrocytes may be responsible for the reduced neuronal activity noted in these cultures. A number of mechanisms by which astrocytes can modulate neuronal activity have been described including modulating calcium signaling, neurotransmitter re-uptake, gliotransmitter release, synapse formation and pruning^52,53^. In principle these could be altered in OCRL depleted cultures leading to a non-cell autonomous impact on neuronal activity. Future experiments will be needed to address such mechanisms.

Why do LS patient derived cultures show an enhanced number of GFAP positive astrocytes during development *in vitro*? Through a single cell multiome analysis of NSC, we found that compared to controls, NSC derived from LS patients contained a larger proportion of cells with transcripts and signatures of astrocytes and glioblasts than neurons when compared to controls (Fig 4). This finding suggests that in LS cultures loss of OCRL function may lead to a developmental switch that results in an increased number of glial precursor cells which in turn leads to increased levels of GFAP noted when these are differentiated.

The ratio of neurons to glia cells in any brain is precisely set in animal species^54^. Both neurons and glia are specified during development through signaling mechanisms including the evolutionarily conserved Notch signaling pathway^55,56^. During our analysis we found multiple lines of evidence supporting elevated Notch signaling in *OCRL* depleted NSC including enhanced levels of cNotch protein and elevation of cNotch dependent transcripts such as *Hes5*. These observations suggest that OCRL depletion results in enhanced Notch signaling thus altering cell specification during brain development. Previous studies have shown that the cleavage of Notch by the γ-secretase enzyme is a process influenced by PIP_2_, thus providing a molecular link between the elevated cNotch levels and the enhanced PIP_2_ in OCRL depleted cells^57–59^. Our finding that levels of the atypical Notch ligand DLK1 were elevated in LSP NSC and secreted DLK1 in 3D organoid cultures provide an additional mechanism for enhanced Notch signaling. Together these finding suggest that during the developmental transition from pluripotent stem cells to NSC, *OCRL* depletion leads to enhanced Notch signaling. Therefore, this may be the reason why NSC derived from OCRL depleted stem cells show enhanced levels of gliogenic precursors. In mouse models, loss of DLK1 shows a dose dependent effect on hippocampal neurogenesis and these mice show defects in cognitive behaviour^41,60^.

Overall, our study provides insights into the cellular and developmental mechanism by which OCRL loss in LS patients may lead to cognitive deficits evident at birth and during early childhood. Our description of a cellular phenotype in a “disease-in-a-dish” model along with the finding the pharmacological inhibition of PIP5KC can reverse this phenotype opens avenues for screening for compounds, especially those with prior FDA approvals for those that can reverse the cellular phenotypes^61^. Such an approach can rapidly accelerate the development of suitable therapeutics for clinical management of LS patients.

## Supporting information

Combined supplementary Figures

## Acknowledgements

This work was supported by the Department of Atomic Energy, Government of India, under Project Identification No. RTI 4006, a Wellcome-DBT India Alliance Senior Fellowship to PR (IA/S/14/2/501540), Department of Biotechnology, Government of India (BT/PR17316/MED/31/326/2015), the Pratiksha Trust and Rohini Nilekani Philanthropies. We thank the NCBS Imaging, Mass spectrometry, Stem Cell and NGS sequencing facility for support. We thank Zubin Rashid for advice on electrophysiology recordings and numerous colleagues for insightful discussions on this study.

## Materials and Methods

### Antibodies

| Name | Catalog | Dilution | Species |
| --- | --- | --- | --- |
| Nestin | ab22035 | 1:200 | Mouse |
| Pax6 | 901301 | 1:200 | Rabbit |
| Foxg1 | ab18259 | 1:200 | Rabbit |
| GABA | A2052 | 1:100 | Rabbit |
| CTIP2 | ab18465 | 1:800 | Rat |
| GFAP | NBP1-05198 | 1:1000 | Chicken |
| S100β | ab52642 | 1:1000 | Rabbit |
| Cleaved-Notch | CST- 4147S | 1:500 | Rabbit |
| DLK1 | ab21682 | 1:500 | Rabbit |
| GAPDH | NB100-56875 | 1:1000 | Rabbit |
| OCRL | sc 393577 | 1:300 | Mouse |
| TBR2 | ab23345 | 1:200 | Rabbit |
| Synapsin-1 | ab64581 | 1:1000 | Rabbit |
| Ki-67 | CST-9449S | 1:100 | Mouse |
| MAP2 | NB300-213 | 1:1000 | Chicken |

List of *secondary antibodies* used
| Name | Catalog | Dilution | Species |
| --- | --- | --- | --- |
| Alexa Fluor™ 488 | A11034 | 1:300 | Goat α-Rabbit IgG |
| Alexa Fluor™ 568 | A11041 | 1:300 | Goat α-Chicken IgG |
| Alexa Fluor™ 488 | A11001 | 1:300 | Goat α-Mouse IgG |
| Alexa Fluor™ 488 | A11006 | 1:300 | Goat α-Rat IgG |
| Alexa Fluor™ 568 | A10042 | 1:300 | Donkey α-Rabbit IgG |
| Alexa Fluor™ 568 | A11004 | 1:300 | Goat α-Mouse IgG |

### Maintenance of LS-patient iPSCs

The generation and characterization of LS patient iPSC has been previously described in Akhtar et.al^27^. hiPSC were maintained StemFlex medium (Gibco, # A3349401) and maintained on hESC-qualified Matrigel coated surface (Corning # 354277).. They routinely passaged with EDTA (Sigma #E8008) and frozen stocks were prepared using PSC cryomix (Gibco, #A26444-01). The cultures were regularly checked for mycoplasma using the Lonza mycoplasma detection kit (# LT07-318).

### Generation of OCRL^KO^ iPSCs

The OCRL knockout iPSC line (iPSC-OCRL^KO^) was generated using CRISPR-Cas9 gene editing in the iPSC wildtype cell line NCRM5^8^. Two sgRNAs, OCRL-688-G1 and OCRL-688-G2, were designed to target OCRL exon 8, based on the c.688C>T truncating mutation identified in the LS patients.

### Cloning of sgRNAs

Two sgRNAs (OCRL-688-G1, 5’ – AAACAATACCCAATCTGGGCAGC – 3’ and OCRL-688-G2, 5’ -CACCGGGTCTCATCAAACATATCC - 3’) were designed to target *OCRL* exon 8. The two sgRNA fragments under the control of their independent U6 promoters were cloned into the plasmid PX458, pSpCas9(BB)-2A-GFP, (Addgene #48138) by Gibson assembly. First, the annealed OCRL-688-G1 was ligated into the linearized PX458 vector using the Bbs1 site. The confirmed clone of PX458-OCRL-688-G1 was then digested with Xba1 and Kpn1 to produce a linearised vector for Gibson assembly. For generation of the insert, the U6-OCRL-688-G2 cassette was amplified from a previously generated clone of PX461-OCRL-688-G2 using custom primers (GA_FP, 5’ –ctgcagacaaatggctctagaGAGGGCCTATTTCCCATG- 3’ and GA_RP, 5’ – agttatgtaacgggtaccGCCATTTGTCTGCAGAATTG- 3’) to introduce a spacer into the amplicon. The U6-OCRL-688-G2 cassette was then inserted into the PX458-OCRL-688-G1 vector using NEBuilder HiFi DNA Assembly Cloning Kit (New England Biolabs), according to the manufacturer’s instructions. The resulting plasmid, PX458-OCRL-688-G1-G2, was purified and presence of both sgRNAs in the final CRISPR-Cas9 vector was confirmed by Sanger sequencing.

### Generation of *OCRL* knockout iPSC line

NCRM5 cells (3×10^6^) were transfected with the assembled CRISPR-Cas9 vector, PX458-OCRL-688-G1-G2, (18µg) using the Neon transfection system and Neon transfection 10 µL kit (Invitrogen Cat # MPK1025). Briefly, cells pre-incubated in fresh E8 Complete media with 1x Revita Cell Supplement (Gibco #A26445-01) for 2 hours were harvested by enzymatic dissociation using StemPro Accutase (Gibco, #A11105-01), for 5 min at 37 °C. After neutralising Accutase with the spent media, the cells were spun down at 1200 RPM for 3 min at room temperature. The cells were resuspended in buffer-R such that each 10 µL hit contained 3 µg of the CRISPR-Cas9 vector and 0.5 ×10^6^ cells. The cell suspension was electroplated using the Neon Electroporation System as per manufacturer’s protocol under the following standardized parameters (1100V, 20ms width, 3 pulses) and plated in a hESC-qualified Matrigel matrix (Corning Cat # 354277) coated 35mm plate pre-incubated with fresh E8 Complete media and Revita Cell Supplement. A complete media change was performed 4 hours post electroporation to remove dead cells and debris. After 24 hours, the cells were checked for GFP expression and were subjected to fluorescence-activated cell sorting (FACS) to enrich for GFP-positive cells. The mixed cell culture was enzymatically dissociated using StemPro Accutase, washed once with PBS and resuspended in media with Penicillin Streptomycin at a concentration of 2×10^6^ cells per mL. The cells were sorted using FACS-Aria Fusion (BD Biosciences). Forward and side scatter parameters were adjusted to eliminate cell clumps and debris. The gating parameter threshold for GFP-positive cells was set using control cells electroplated with pCXLE-EGFP (Plasmid #27082). The GFP-positive cells were plated in a pre-incubated Matrigel coated 60 mm dish. Clonal expansion of isolated colonies was performed manually. gDNA was isolated from stable colonies using QIAamp DNA Mini Kit (Qiagen Cat # 51304) and the target region was amplified using specific primers (OCRL_E8_PCR_F, 5’ –TCAAAGCCCTGTTACCCTGG- 3’ and OCRL_E8_PCR_R, 5’ –GACAGGAGCTTGAAACAGGC- 3’). The PCR product was purified and clones with the desired out-of-frame indel modifications were identified by Sanger Sequencing.

### Generation of neural stem cells from iPSCs and terminal differentiation into neurons

Differentiation of forebrain cortical neurons from iPSC was carried out as per the published protocol ^62^Briefly, to generate dorsal forebrain neural stem cells, iPS cells were plated on 6-well tissue culture plates coated with Matrigel.When cells reach 100% confluency, medium was replaced with neural induction (NI) medium marking 0 days *in vitro* (DIV) and maintained for 10 DIV. The NI medium consisted of Neural Maintainence Media (NMM): (1:1) Neurobasal (Gibco, #21103049) and DMEM/F12+Glutamax (Gibco, #10565018), insulin (2.5ug/ml, Sigma #11061-68-0), B27 with vitamin A (1X, Gibco #17504044), N2 supplement (1X, Gibco #17502048), betamercaptoethanol (1X, Gibco #21985-023), sodium pyruvate (1mM, Gibco #11360-070), non-essential amino acids (NEAA, 1X, Gibco #11140-050), penstrep (1000U/ml, Gibco #15140-122), glutamax (1mM, Gibco #35050-061), supplemented with SB-43142 (10uM, Sigma #301836-41-9), and LDN (200nM, Sigma #SML0559). On 10^th^ day, cells were dissociated using dispase (Life Technologies) and resuspended in NI medium with 1X revita supplement. Cells were plated on air-dried poly-L-ornithine (Merck # A-004-C) and 10ug/ml laminin-coated (Gibco # 23017015) in 6-well plates. The following day, NIM was replaced with neural maintenance (NM) medium supplemented with 20 ng/ml FGF2 (Gibco #PHG0369), which was added for 4 days. Lastly, NPCs were frozen at 1 million density in PSC cryomix. For terminal differentiation of NPCs into cortical neurons, ∼400,000–500,000 NPCs were plated in glass bottom dishes and 6-well dishes cotaed with poly-L-ornithine/laminin-coated plates and maintained in NM media supplemented with 20ng/ml BDNF (Gibco # PHC7074), GDNF (Gibco #PHC7041), ascorbic acid (50uM, Sigma # A4544), and dbcAMP (50uM, Sigma #D0627). During the first 14 days after plating, cells were treated with 10 μM DAPT (Sigma-Aldrich #D5942) to synchronize the neuronal maturation process.

### Generation of dorsal forebrain organoids

Brain organoids were generated from iPSC using a modified protocol based on Pasca lab protocol^63^. Briefly, iPSC were dissociated with accutase to obtain a single-cell suspension. Approximately 3 × 10^6 cells were placed in each well of an AggreWell 800 plate (STEMCELL Technologies # 34811) containing Stemflex medium enriched with 1X Revita supplement. The plate underwent centrifugation at 100g for 3 minutes and was then incubated at 37°C with 5% CO_2_.

### Neural induction

After 48 hours (designated as day 0), cell aggregates were transferred to ultra-low attachment dishes (6-well or 60mm and 100mm) and cultured in Essential 6 medium supplemented with LDN (200nM) and SB-431542 (10 μM). The medium remained unchanged on day 1. To prevent spontaneous fusion of EBs, dishes were transferred to shaker from day 0 onwards. From days 2 to 5, organoids were maintained in E6 medium containing patterning molecules.

### Expansion of neural stem cells

On day 6, organoids were moved to a neural medium composed of Neurobasal A (Gibco #10888022), B-27 supplement (vitamin A-free), GlutaMax (1:100), and penicillin-streptomycin (1:100). This medium was further supplemented with EGF (20 ng/ml) and FGF2 (20 ng/ml) from day 6 to 24. Medium changes were performed daily for the initial 10 days, followed by every other day for the subsequent 9 days.

### Neuronal maturation

From day 25 to day 43, neural medium was supplemented with 20ng/ml brain-derived neurotrophic factor (Peprotech, #450-02) and 20ng/ml NT-3 (Peprotech, #450-03). From day 44, only neural medium without growth factors was used for medium changes every 4 days and B27 (vitamin A-free) was replaced with B27 plus supplement (Gibco #A3582801).

### Cryopreservation, sectioning, IHC of organoids and ICC for monolayer cultures

Organoids were fixed in 4% PFA-PBS overnight at 4 °C followed by dehydration in 30% sucrose for 24-48hrs. Subsequently, samples were embedded in optimal cutting temperature (OCT) compound (Tissue-Tek OCT Compound 4583, Sakura Finetek) and 30% sucrose–PBS (1:1) for cryo-sectioning (20-μm-thick sections) using a Leica Cryostat (Leica, catalogue no. CM1860). For immunofluorescence staining, cryosections were washed with PBS to remove excess OCT. Cryosections were blocked in 5% BSA and 0.3% Triton X-100 diluted in PBS for 1 h at room temperature. Sections were then incubated overnight at 4 °C with primary antibodies diluted in PBS containing 5% BSA and 0.1% Triton X-100. PBS was used to wash away excess primary antibodies, and the cryosections were incubated with secondary antibodies in PBS containing 5% BSA for 1 h. The nuclei were visualized with Hoechst33258 (Thermo Fisher Science, catalogue no. H3569, 1:10,000 dilution).

NSC and neuronal cultures were fixed using 4% formaldehyde in phosphate-buffered saline (PBS) for 30 min and permeabilized using 0.1% TX-100 for 5 min. These cultures were incubated at room temperature for 1 h in a blocking solution of 5% BSA in PBS. Primary antibodies at respective dilutions were added and incubated overnight at 4°C in blocking solution, followed by incubation with secondary antibodies in blocking solution (Invitrogen) for 1 h. Antibodies used were as follows: Nestin (abcam) 1:200; Pax6 (abcam) 1:200; DCX (abcam) 1:200; SATB2 (abcam) 1:300; MAP2 (abcam) 1:1000; GFAP (abcam) 1:100, S100B(abcam) 1:500; TBR1 (abcam) 1:300; FOXG1 (abcam) 1:200; OCRL (C-terminus, Santacruz) 1:300; OCRL (N-terminus, Sigma) 1:300. Confocal images were recorded by collecting a range of z-stack and the image stack was merged using Z-project (maximum intensity projection) function using ImageJ (National Institute of Health, USA, http://imagej.nih.gov/ij).

### Calcium imaging

Calcium imaging was according to our previously published protocol with minor modifications ^64^. Briefly, neurons were washed with Tyrode’s buffer solution (5 mM KCl, 129 mM NaCl, 2 mM CaCl_2_, 1 mM MgCl_2_, 30 mM glucose and 25 mM HEPES, pH 7.4) for 10 min and later incubated with 4 uM fluo-4/AM (1 mM, Molecular probes, #F14201) and 0.02% pluronic F-127 (Sigma-Aldrich, #P2443) in the Tyrode’s buffer solution in the dark for 30–45 min at room temperature. Following dye loading, the cells were washed again with the buffer thrice, each wash for 5 min. Finally, cells were incubated for an additional 20 min at room temperature to facilitate de-esterification. Ca^2+^ imaging was performed for 10 min with a time interval of 1s at 20X objective of wide-field fluorescence microscope Olympus IX-83. A 4-min baseline measurement was recorded to visualize calcium transients, followed by the addition of TTX to abolish calcium transients for another 4 min, and finally high KCL was added as an internal control. Calcium traces were obtained using the Imagej software by drawing a region of interest (ROI) manually around each neuronal soma. To plot the calcium traces, the raw fluorescence intensity values from each neuron were normalized to the first fluorescence intensity signal of the baseline recording. Frequency of calcium transients was measured.

### Electrophysiology

Whole-cell patch clamp recordings were performed on 60DIV iPSC-derived cortical neurons. Recordings were performed at room temperature. Briefly, coverslips containing the cultures were transferred to a recording chamber perfused with an external solution (in mM: 152 NaCl, 2.8 KCl, 10 HEPES, 2 CaCl_2_, 10 glucose; pH 7.3–7.4, 300–320 mOsm). A MultiClamp 700B amplifier was used to collect recordings, with data sampled at 10 kHz and digitized at 20 kHz via a Digidata 1550. Patch pipettes were crafted from thick-walled borosilicate glass and filled with an internal solution (in mM: 155 K-gluconate, 2 MgCl_2_, 10 HEPES, 10 Na-PiCreatine, 2 Mg^2+^-ATP, 0.3 Na3-GTP; pH 7.3, 280–290 mOsm). Series resistance was typically 5–8 MΩ. Membrane voltage was held at -70 mV. Currents were recorded using a voltage-step protocol. The membrane potential was initially held at -20 mV and then incrementally increased in 5 mV steps up to +100 mV. Stimulation protocols were designed using pClamp 10.5 software, with subsequent analysis performed offline using Clampfit 11.1 software.

### Western blotting

Cultures were harvested using Stempro Accutase and pelleted at 1000 g for 5 min then washed three times with ice-cold PBS. The pelleted cells were homogenized in 1X RIPA lysis buffer containing freshly added phosphatase and protease inhibitor cocktail (Roche). To remove cellular debris, crude RIPA lysates were centrifuged at 13,000 rpm for 20 min at 4°C. The supernatant was transferred to a new tube and quantified with a Pierce BCA protein assay (Thermo Fisher Scientific, #23225). Thereafter, the samples were heated at 95°C with Laemmli loading buffer for 5 min and 20 ug protein was loaded onto Bolt™ 4 to 12%, Bis-Tris SDS gel (Invitrogen, #NW04120BOX). The proteins were then transferred onto a nitrocellulose membrane and incubated overnight at 4°C with indicated antibodies. The blots were then washed three times with Tris Buffer Saline containing 0.1% Tween-20 (0.1% TBS-T) and incubated with 1:10,000 concentration of appropriate HRP-conjugated secondary antibodies (Jackson Laboratories, Inc.) for 45 min. After three washes with 0.1% TBS-T, blots were developed using Clarity Western ECL substrate (Bio-Rad) on a GE ImageQuant LAS 4000 system.

### Mass spectrometry analysis

Lipid extraction followed by liquid chromatography-mass spectrometry was used to estimate PIP_2_ levels. The methods used are identical to that reported in Akhtar et.al.^27^

### 10x Multiomics

#### Nuclei isolation from NSC

NSC were revived and nuclei were isolated as per 10x genomics recommended protocol. Briefly, approximately 1.2×10^6^ NSCs were thawed, resuspended in PBS + 0.04% BSA and spun down at 1250rpm for 5mins. The cell pellet was resuspended in lysis buffer containing Tris-HCl (pH 7.4) 10mM, NaCl 10mM, MgCl_2_ 3mM, Tween-20 0.1%, IGEPAL CA-630 0.1%, Digitonin 0.01%, BSA 1%, DTT 1mM, RNase inhibitor (Roche) 1U/ul. The cell pellet was allowed to lyse for a period 3mins. Chilled wash buffer containing Tris-HCl (pH 7.4) 10mM, NaCl 10mM, MgCl_2_ 3mM, BSA 1%, Tween-20 0.1%, DTT 1mM, RNase inhibitor 1U/ul was added to lysed cells and centrifuged 1250rpm for 5mins at 4°C. The cells were washed for two more times. Finally, the cells were resuspended in nuclei buffer containing DPBS with DTT 1mM, and RNase inhibitor 1 U/μl. Nuclei were counted using a hemocytometer, diluted to 10000 nuclei, and further processed following 10x Genomics Chromium Next GEM Single Cell Multiome ATAC + Gene Expression Reagent Kits user guide. We targeted 10,000 nuclei per sample per reaction. Libraries from individual samples were pooled and sequenced on the NovaSeq 6000 sequencing system, targeting 2×50 base pairs reads for ATAC as well as RNA.

#### Pre-processing of snMultiome data

The raw sequencing signals generated from NovaSeq 6000 in the BCL format were demultiplexed into fastq format using the “mkfastq” function in the Cell Ranger ARC suite (10x Genomics). Cell Ranger-ARC count pipeline was implemented for cell barcode calling, read alignment, and quality assessment using the human reference genome (GRCh38) following the protocols described by 10x Genomics. The final outs file so generated for each sample were used for downstream analysis using Scanpy for RNA and SnapATAC2 and pycisTopic for ATAC.

#### RNAseq pre-processing

The resulting matrix from the Cell Ranger ARC pipeline was imported into Scanpy using the scanpy.read_10x_mtx() function for WT1, LSP2 and LSP3 NSCs. The three samples were concatenated and used for further processing. Quality control metrics were calculated for each nucleus, including the number of genes expressed, total count of unique molecular identifiers (UMIs), and the percentage of mitochondrial gene expression. After initial preprocessing, we performed quality control and filtering steps to remove low-quality nuclei and genes with low detection rates. This process was implemented using Scanpy’s preprocessing functions. Specifically, we applied the following filters: we removed nuclei expressing fewer than 200 genes using the sc.pp.filter_cells() function and genes detected in fewer than 5 nuclei were filtered out using the sc.pp.filter_genes() function. Potential doublets were identified using Scrublet, implemented via sc.pp.scrublet(), with sample information provided as a batch key to account for sample-specific doublet rates. Data was then normalized to the median of total counts across all cells using sc.pp.normalize_total(). A log-transformation (log(1+x)) was applied using sc.pp.log1p() to account for the large dynamic range of expression values. Highly variable genes were identified using sc.pp.highly_variable_genes(), selecting the top 2000 genes across all samples. Principal Component Analysis (PCA) was performed using sc.tl.pca(). A neighborhood graph was constructed using sc.pp.neighbors() based on the PCA representation. Uniform Manifold Approximation and Projection (UMAP) was applied for visualization using sc.tl.umap(). Cells were clustered using the Leiden algorithm, implemented with sc.tl.leiden(). To identify cluster-specific marker genes, we performed differential expression analysis using the Wilcoxon rank-sum test, implemented with sc.tl.rank_genes_groups(). The Leiden based clustering obtained was then used for cell-state annotation using snapseed and scVI-scanVI framework below.

#### Cell-state annotation

We used two ways to annotate cells in our dataset. First, we used Snapseed that uses a pre-defined marker list to provide annotation results. To this end, we used the cell-type marker list from Braun et.al., 2023 and filtered other regions to retain only forebrain, which resulted in neuronal IPC, radial-glia, glioblast, and neuron cell-types. This list was converted to .yaml file as per Snapseed instructions. Second, we employed the scvi-scanvi framework for cell-type annotation of our single-cell RNA sequencing (scRNA-seq) data, to leverage both unsupervised and supervised learning techniques to integrate reference and query datasets and transfer cell-type labels. We first prepared the reference datasets Braun et.al., and Wang et.al. 2024 using scanpy. Both the datasets were filtered to include specific cell populations of interest. Specifically, for Wang.et.al., we retained cells from first, second, and third trimesters, and focusing on cells identified in their ‘subclass’ category: Astrocytes, Cajal-Retzius cell, GABAergic neuron, Glutamatergic neuron, IPC-EN, IPC-Glia, OPC, Oligodendrocyte and Radial-glia. To ease computing resource, we down sampled the dataset to 80,000 cells for Wang et.al., and ∼2×10^5^ cells for Braun et.al. dataset. Quality control was performed by filtering out cells with fewer than 300 expressed genes. Highly variable genes (HVGs) were identified using Scanpy’s highly_variable_genes function, selecting the top 2,000 genes. We employed the SCVI (Single-Cell Variational Inference) model for dimensionality reduction and data integration. The data was log-normalized, and the SCVI model was set up and trained on the reference dataset. Latent representations were extracted for downstream analysis. Using the SCVI latent representations, we constructed a neighbor graph and performed Leiden clustering for unsupervised cell grouping. The cell type labels were prepared for supervised learning using the SCANVI model. The WT1, LSP2, LSP3 NSCs processed query dataset was loaded and preprocessed similarly to the reference dataset as outlined. We identified common genes between the reference and query datasets, subsetting both to these shared highly variable genes to ensure compatibility for integration. For dataset integration and cell type prediction, we employed the SCANVI (Single-Cell Annotation using Variational Inference) model. The model was trained on the reference dataset and then applied to the query dataset. SCANVI latent representations were extracted, and cell types were predicted for the query cells based on the reference dataset annotations. We selected highly confident predictions for Braun et.al.^32^, followed by Wang et.al.^23^, through the confusion matrix. Finally, based on the consensus from Snapseed, scVI-scanVI framework, the 14 cell clusters in our NSC-dataset were labelled.

#### SCENIC+ analysis

We employed the SCENIC+ workflow to construct gene regulatory networks for the clusters in our NSC dataset. The analysis pipeline began with consensus peak calling for each cell type using MACS2 with the default parameters. We implemented the Latent Dirichlet Allocation (LDA) algorithm using MALLET. The default parameters were used, with 500 iterations for topic modeling. Topics were binarized using the Otsu method to convert continuous distributions into discrete region sets. Both differentially accessible regions (DARs) and topics (sets of co-accessible regions) across cell types were used as candidate enhancers. Outputs of pycisTopic cistopic object and scanpy were used as input for SCENIC+ analysis as outlined in their tutorial default settings^33^.

#### snapATAC2

ATAC-seq data was imported using SnapATAC2’s import_data function, utilizing the hg38 genome as reference. Initial quality control metrics, including the number of fragments, fraction of duplicates, and fraction of mitochondrial reads, were analyzed with default settings as per the tutorial. Cells were filtered based on the number of counts (5,000-100,000) and minimum TSSE score (>10). A tile matrix was generated, and 250,000 features were selected for downstream analysis. Potential doublets were identified and filtered using the Scrublet algorithm. Dimensionality reduction was performed using spectral decomposition, followed by k-nearest neighbor graph construction and Leiden clustering for initial cell grouping. A gene activity matrix was created using the make_gene_matrix function, associating ATAC-seq peaks with gene annotations from the hg38 genome.

To annotate and compare ATAC cell clusters, we integrated our ATAC-seq data with its paired RNA NSC dataset annotated above. The query (ATAC-seq) and reference (RNA-seq) datasets were concatenated, retaining only genes present in both datasets. Highly variable genes (top 5,000) were identified using Seurat v3 methodology. We utilized the scvi-tools package for data integration and cell type prediction. A Variational Autoencoder (VAE) model was trained on the combined dataset using the SCVI class, with 2 layers, 30 latent dimensions, and negative binomial gene likelihood. The model was trained for up to 1,000 epochs with early stopping. Subsequently, a semi-supervised SCANVI model was initialized from the trained SCVI model. Known cell type labels from the reference dataset were used to guide the annotation of query cells. The SCANVI model was trained for up to 1,000 epochs, sampling 100 cells per label in each epoch. Cell type predictions for the query dataset were obtained using the trained SCANVI model. The latent representations from SCANVI were used for neighbor graph construction and UMAP visualization. The predicted cell types were then mapped back to the original ATAC-seq object.

#### RNA isolation, cDNA conversion and quantitative PCR

RNA from three biological replicates of 2D NSCs/neurons and 3D organoids were extracted using TRIzol (Ambion, Life Technologies), as per manufacturer’s instructions. The total RNA was then quantified using NanoDrop 1,000 spectrophotometer (Thermo Fisher Scientific). 1 μg of RNA was used for DNase I treatment in a reaction mixture of 10 mM DTT and 40U RNase inhibitor (RNaseOUT, Thermo Fisher Scientific). This reaction mixture was then incubated at 37°C for 30 minutes followed by 70°C heat inactivation for 10 minutes. For cDNA synthesis, 200U of Superscript II Reverse Transcriptase (Invitrogen) was added to the reaction mixture along with 2.5 μM random hexamers and 0.5 mM dNTPs. The reaction mixture was then incubated at 25°C for 10 minutes, followed by 42°C for 60 minutes and heat inactivated at 70°C for 10 minutes, along with a no reverse-transcriptase control sample. Real-time quantitative PCR of the cDNA samples was performed using primers for genes of interest and control gene, GAPDH, on Applied Biosystems 7500 fast qRT-PCR system. The reporter used here is Power SYBR Green Master mix (Applied Biosystems). The reaction was run for 40 cycles of 95°C for 30 seconds (denaturation), 60°C for 30 seconds (annealing) and 72°C for 45 seconds (extension). The relative mRNA expression of different genes was then calculated by ΔC_t_ method, normalizing their 2^-ΔCt^ values to GAPDH. Unpaired student t-test with welch correction was used for statistical analysis and p<0.05 was considered significant.

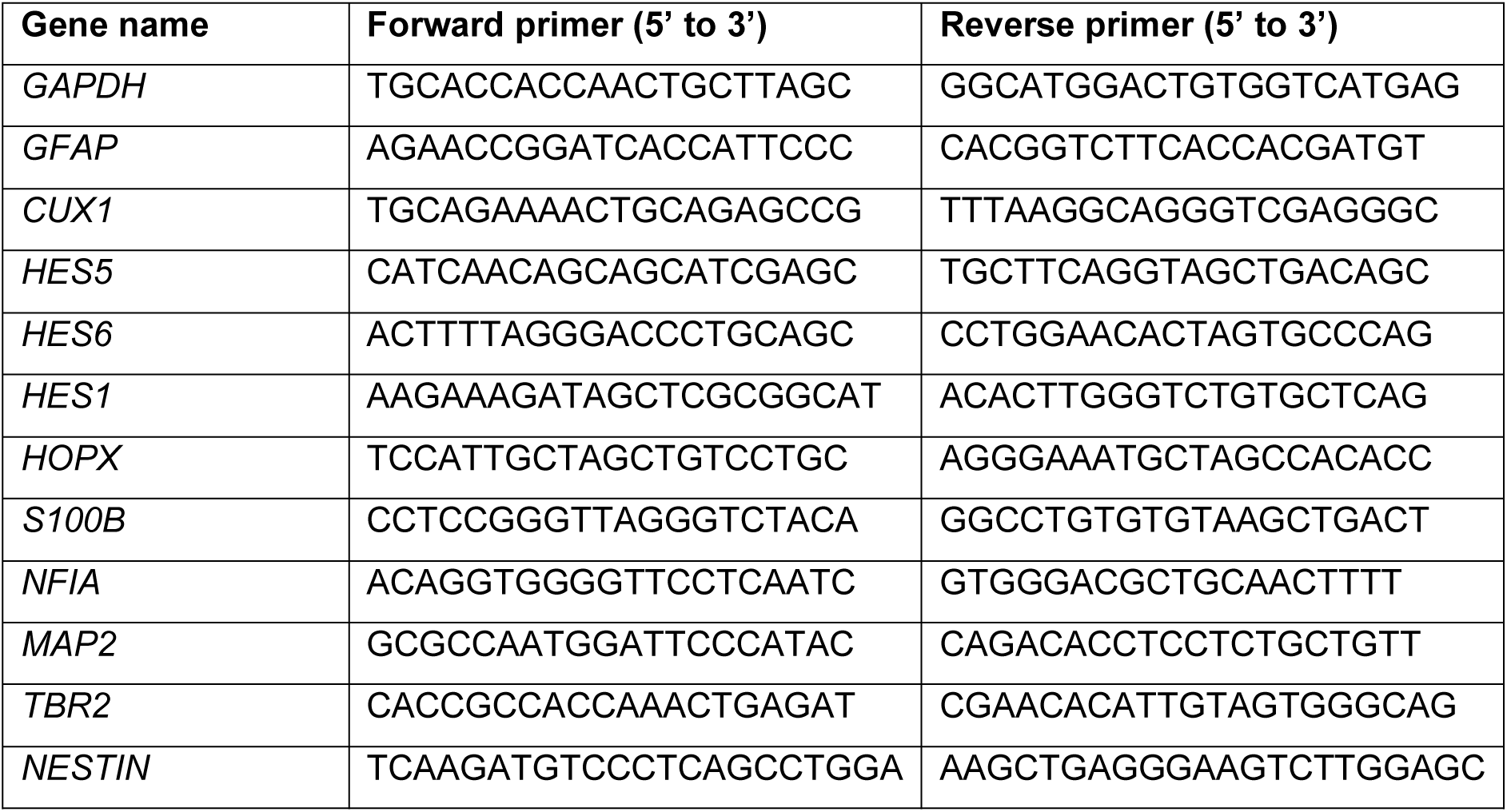

## Supplementary Figure Legends

**Suppl. Figure 1: Generation of OCRL^KO^ cell line**

**(A)** iPSC-derived NSCs using dual-SMAD inhibition exhibit canonical NSC markers Nestin (red), Pax6 and FOXG1 (green). NSCs were counter-stained with DAPI, Scale bar = 50 µm. Immunofluorescence images obtained from WT1, LSP2, LSP3 and LSP4 **(B)** CRISPR-Cas9 targeting strategy: Two sgRNAs (G1 and G2) were designed to target exon 8 of *OCRL*, just prior to the 5’-Phosphatase domain of the protein. Target sites of *OCRL*-688-G1 and *OCRL*-688-G2 are highlighted in yellow and green respectively **(C)** Sequence chromatogram confirming insertion of 11bp (red outlined box) in OCRL^KO^ iPSC (lower panel), compared to non-edited WT2 iPSC (upper panel). **(D)** iPSC from WT2 and OCRL^KO^ displaying pluripotent markers viz., SOX2 (green), SSEA4 (red), Oct-4 (green), TRA-160 (red); iPSCs were counter-stained with DAPI (blue), scale bar=50μm. **(E)** Normal karyogram confirming chromosomal integrity of OCRL^KO^ iPSC **(F)** iPSC-derived NSC from WT2 and OCRL^KO^ exhibit canonical NSC markers Nestin (red), Pax6 and FOXG1 (green). NSCs were counter-stained with DAPI, scale bar=50μm.

**Suppl. Figure 2: Physiological properties of iPSC derived neurons**

**(A)** Resting membrane potential from WT2 and OCRL^KO^ derived 60 DIV neurons measured during whole cell recordings. Y-axis show resting membrane potential in mV. Each point represents a single cell. Mean +/- SEM shown. **(B)** Capacitance measurements from WT2 and OCRL^KO^ 60 DIV neurons measured during whole cell recordings. Y-axis shows capacitance in pF. Each point represents a single cell. Mean +/- EM shown. Whole-cell patch clamp recordings in voltage-clamp mode from 60DIV WT1 and pooled LSP neurons **showing (C)** inward and **(D)** outward currents. Y-axis shows currents in pA. X-axis voltage in mV. Averaged traces from WT1 (11 cells) and LSP (6 cells) is shown. Current amplitude at each voltage is shown as mean +/- SEM.

**Suppl Figure 3: Overview of single nuclei multiome data analysis**

**(A)** Diagram illustrating the workflow and steps for generating and analysing single nuclei multiome data. **(B)** UMAP projection showing cell clustering based on the Leiden algorithm using the scanpy pipeline. Each colour represents a distinct cell cluster. **(C)** UMAP coloured by cell class, highlighting distribution of major cell types such as glioblasts, neuroblasts, neurons, neuronal IPCs, and radial glia; data set from Braun et.al ^32^. **(D)** Confusion matrix from scVI-scANVI mapping; Y-axis depicts the 14 unannotated clusters obtained from our multiome analysis; X-axis are the cell types seen in the Braun et.al dataset showing prediction accuracy for different cell types across conditions. **(E)** Confusion matrix from scVI-scANVI mapping, detailing prediction outcomes for various cell clusters in Wang.et.al. 2024^23^, dataset. **(F)** Dot plot displaying expression levels of marker genes across cell clusters generated using scanpy. Dot size indicates the percentage of cells expressing each gene, while colour intensity reflects expression level. Upper X-axis shows cell clusters numbered 0-13; Lower X-axis included names of transcripts whose enrichment is being plotted. Y-axis show annotated cell clusters **(G)** UMAP projection showing sample distribution (WT1, LSP2, LSP3) using the snapATAC2 pipeline. **(H)** UMAP projection with Leiden clustering across all samples, indicating distinct clusters identified through the snapATAC2 analysis.

**Suppl Figure 4: 3D spheroids generated from iPSC**

Phase contrast images showing the morphology of WT1, LSP2, LSP3, LSP4 **(A)** and WT2 and OCRL^KO^ **(B)** neural spheroids across specific developmental timepoints: 6, 15, 30 and 60 DIV. Scale bar: 500μm RT-PCR analysis of spheroids derived from the LSP iPSC lines compared to WT1. Transcript levels for the neuronal marker MAP2 **(C)** NF1A **(D)** and GFAP **(E)** are shown. Y-axis depicts transcript levels normalised to the housekeeping gene GAPDH. X-axis shows genotypes. Each point depicts transcripts measured from extracts of 15-20 spheroids. Error bars mean +/- SEM. Statistical test: Unpaired t-test with Welch correction. **(F)** Confocal images of 90 DIV organoids from WT1, LSP2, LSP3 and LSP4. Scale bar: 50μm. GFAP (red) S100β (green) DAPI (blue).

## Supplementary Table legends

**Suppl Table 1: Transcription factor motif analysis from multiome data**

The table describes enriched motifs for each cell cluster obtained through SCENIC+ analysis. Motif logo shows y-axis representing the information content measured in bits, ranging from 0 to 2 bits for DNA sequences. Specifically, a value of 0 bits indicates a position where all nucleotides occur with equal probability. A value of 2 bits indicates a position where only a single nucleotide occurs. The height of each stack shows how conserved that position is - taller stacks mean higher conservation. The x-axis shows the nucleotide positions in the DNA sequence alignment. Each position contains a stack of letters (A, C, G, T) where: the letters are stacked according to their relative frequency at that position. The height of each individual letter within a stack represents how frequently that nucleotide appears at that position and the most common base appears as the largest letter at the top of each stack. In summary, In the motif logo, the x-axis represents sequential nucleotide positions (5’ to 3’), and the y-axis shows information content in bits (0-2.0). Letter height indicates the relative frequency of each nucleotide (A, T, C, G) at each position, with taller letters representing more frequently occurring bases.

