## Supplementary material for "Enhanced Notch dependent gliogenesis and delayed physiological maturation underlie neurodevelopmental defects in Lowe syndrome": Combined supplementary Figures

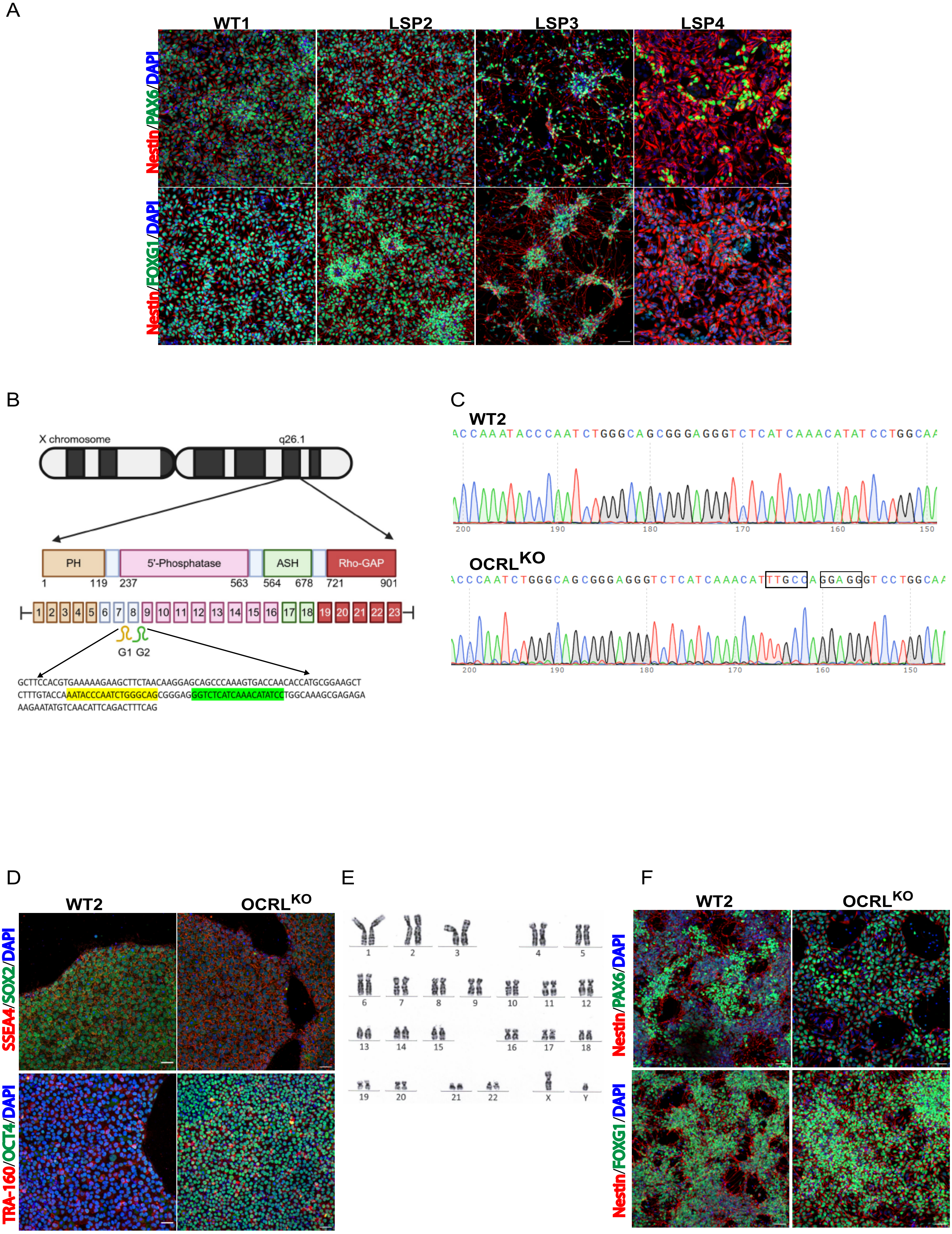


Sharma et.al., Supplementary Fig1

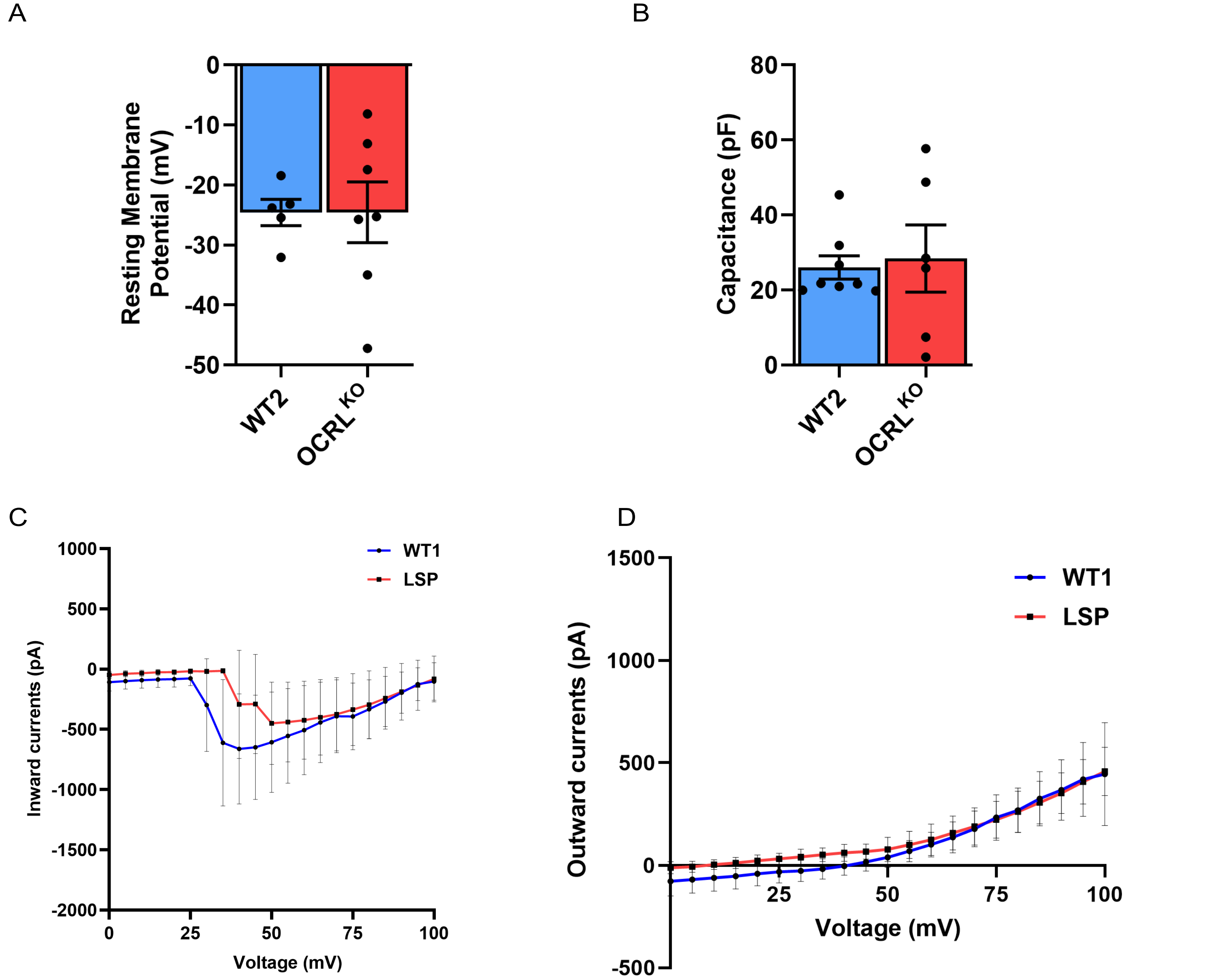


Sharma et.al., Supplementary Fig2

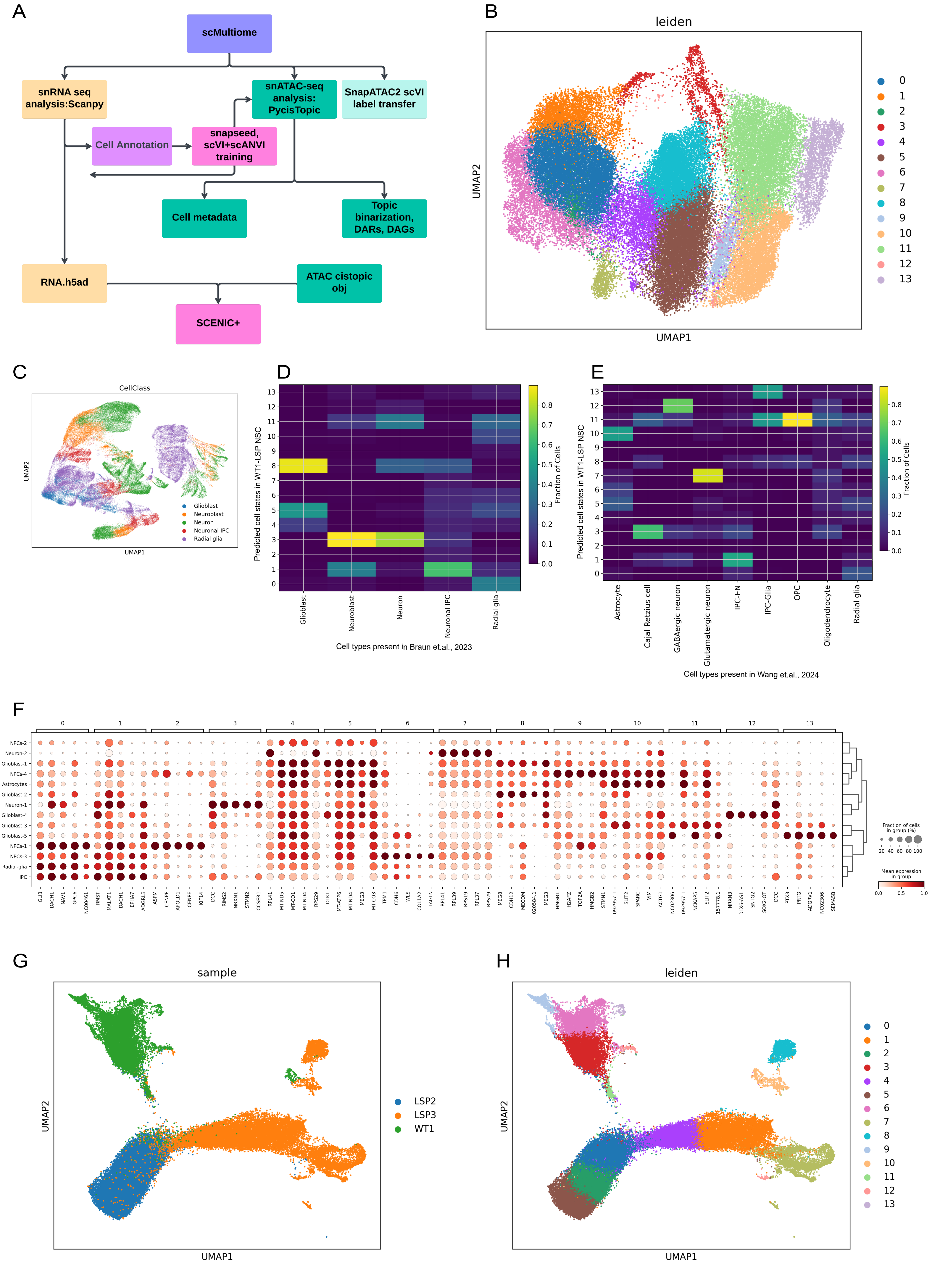


**Sharma et.al., Supplementary Fig3**

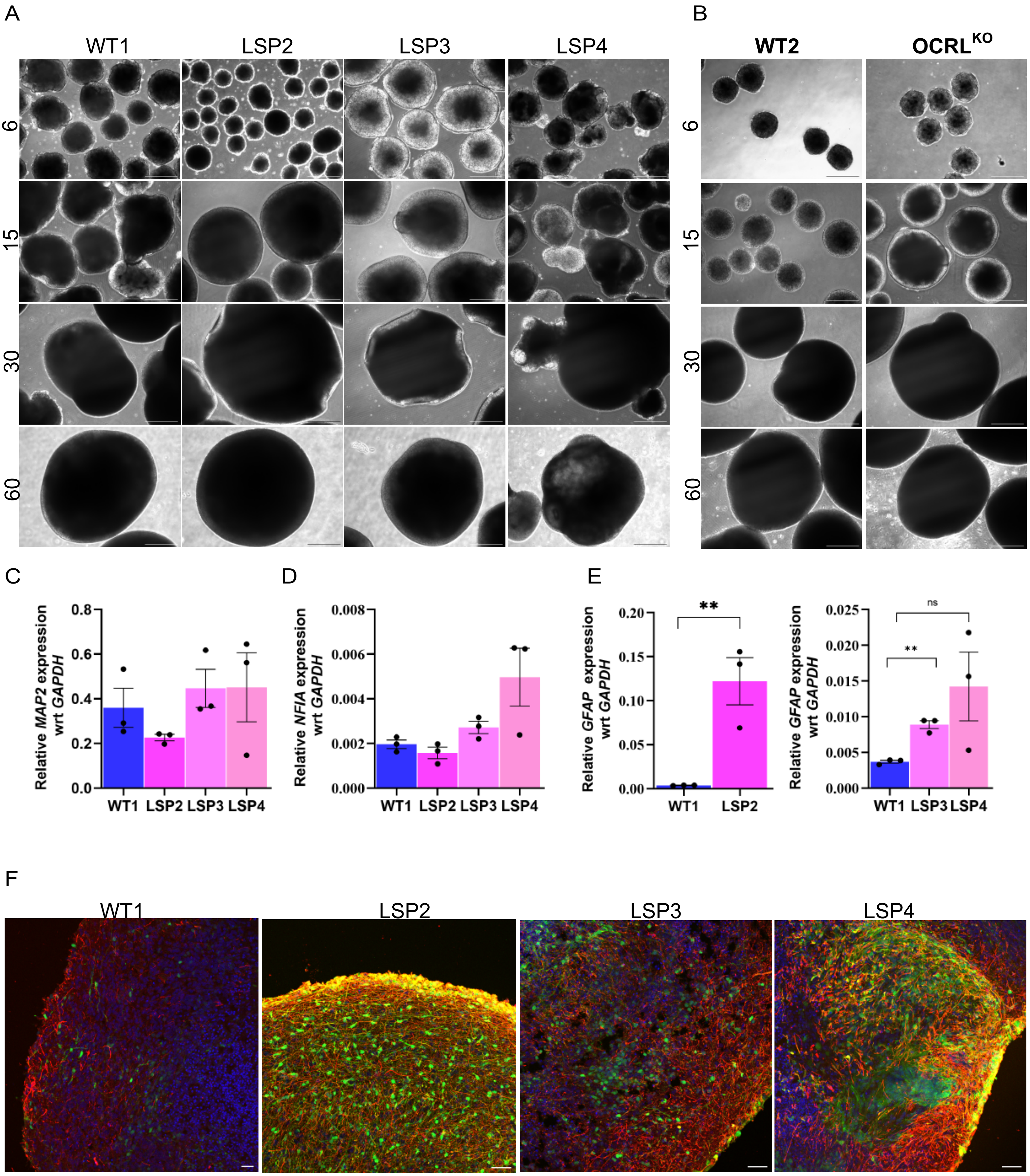


**Sharma et.al., Supplementary Fig4**
