## Supplementary Table for "Enhanced Notch dependent gliogenesis and delayed physiological maturation underlie neurodevelopmental defects in Lowe syndrome"

| Motif logo | TF | Cell-cluster | NES | AUC |
|---|---|---|---|---|
| 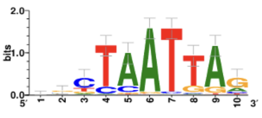 | LHX9 | Radial-glia | 7.269415 | 0.015211 |
| 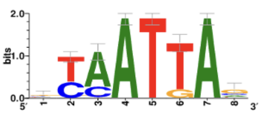 | EMX1 | Radial-glia | 6.380436 | 0.013670 |
| 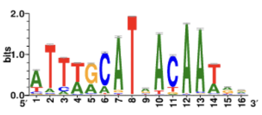 | SOX2, POU2F1 | Glioblast | 7.025670 | 0.013287 |
| 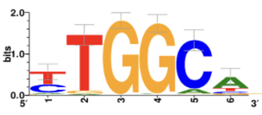 | NFIX, NFIC, NFIB, NFIA | Astrocytes | 6.759462 | 0.012162 |
| 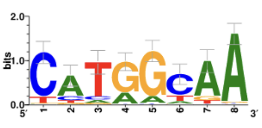 | RFX2, RFX3, RFX4, RFX5, RFX1 | Astrocytes | 18.842255 | 0.028898 |

**Sharma et.al., Supplementary table 1**
